# FOCUS-3D: Robust, generalizable volumetric cell segmentation for three-dimensional fluorescence microscopy

**DOI:** 10.64898/2026.08.25.746907

**Authors:** Qinghua Zhang, Zeyu Mu, Boqi Liu, Yunfeng Chi, Donglin Li, Wenjuan Wang, Jian-Quan Ni, Yinan Wan, Li Yu, Joaquín Navajas Acedo, Guoqiang Yu

## Abstract

Understanding how cells establish spatial organization within tissues is a fundamental question in life sciences. While modern three-dimensional fluorescence microscopy captures large-volume tissue architecture, extracting quantitative cellular insights from complex volumetric datasets remains a major barrier. Here, we introduce FOCUS-3D, a robust, broadly generalizable volumetric cell segmentation framework built on a large, diverse manually annotated cell resource and advanced AI designs. Integrating volumetric representation learning, multi-scale feature extraction, and query-based mask prediction, FOCUS-3D achieves state-of-the-art performance across diverse species, tissues, fluorescent reporters and imaging modalities. During zebrafish (*Danio rerio*) development, FOCUS-3D uncovers three successive phases of notochord morphogenesis. We disentangle early motility-driven rearrangements from later cell shape remodeling and tissue repacking, and further link these morphological states to spatial and developmental transcriptional programs across independent datasets.

## Introduction

Understanding how cells are spatially organized, coordinated, and remodeled within tissues is central to uncovering how biological form and function emerge^1–5^. Advances in three-dimensional (3D) fluorescence microscopy now enable observation of large cell populations in intact embryos, organs, and living tissues at unprecedented spatial and temporal resolution^5,6^, connecting the spatiotemporal dynamic behavior of individual cells to the organization of entire tissues. Yet, these large and complex 3D datasets must be transformed into quantitative, high-quality cellular representations to study tissue organization and dynamics. 3D cell instance segmentation is fundamental to this transformation, so that each cell can be identified within its native spatial context, enabling measurements of cell position, morphology, movement, neighborhood relationships, and collective tissue rearrangement^3,4,7,8^.

Recent innovations in artificial intelligence have greatly advanced cell segmentation^9–11^, yielding a suite of widely used analytical tools^12–20^. However, these approaches are predominantly designed for two-dimensional (2D) imaging and adapted to volumetric data via slice-wise inference, orthogonal-view fusion, or post-hoc stitching and linking^13–17^. Although convenient, such indirect 3D paradigms severely compromise segmentation fidelity by disrupting spatial continuity and distorting volumetric shape information, particularly in densely packed tissues, anisotropic volumes, and morphologically heterogeneous cellular samples^21,22^. Progress toward robust and generalizable native 3D segmentation models is hampered by the scarcity of diverse, well-annotated 3D datasets^23,24^. Creating volumetric instance labels is laborious and requires substantially more time and expertise compared with 2D cell annotation^23,25^. As a result, existing 3D models such as StarDist^26^ were typically trained on narrow biological systems and fail to generalize across species, tissues, reporter strategies, microscopes, and experimental conditions. The absence of unified, broadly generalizable volumetric cell-resolved analysis constitutes a major bottleneck for extracting biological insights from large-scale volumetric imaging data^27,28^.

To address these bottlenecks, we developed FOCUS-3D (<u>F</u>oundational <u>O</u>pen-source <u>C</u>ellular <u>U</u>nified <u>S</u>egmentation across 3D microscopy), a versatile framework for generalizable three-dimensional cell instance segmentation and cell-resolved quantitative analysis (Figure 1). We first constructed what is, to our knowledge, the largest expert-annotated resource for volumetric cell segmentation, containing more than 410,000 manually curated 3D cell instances spanning diverse biological specimens and imaging conditions. We further designed a unified native 3D segmentation model that learns holistic representations from intact microscopy volumes and predicts complete cell instances directly in 3D space, avoiding the inherent limitations of slice-based, 2D decomposition strategies. To fully and efficiently exploit limited manual annotations, FOCUS-3D adopts an integrative multi-tier training paradigm that hierarchically leverages large-scale unlabeled volumes, automatically generated annotations, and high-quality expert-curated labels. By synergistically integrating volumetric representation learning, multi-scale spatial encoding, and query-based instance prediction, our framework captures both fine-grained cellular boundaries and global tissue contextual features^29,30^ (Figure 2).

**Figure 1:**
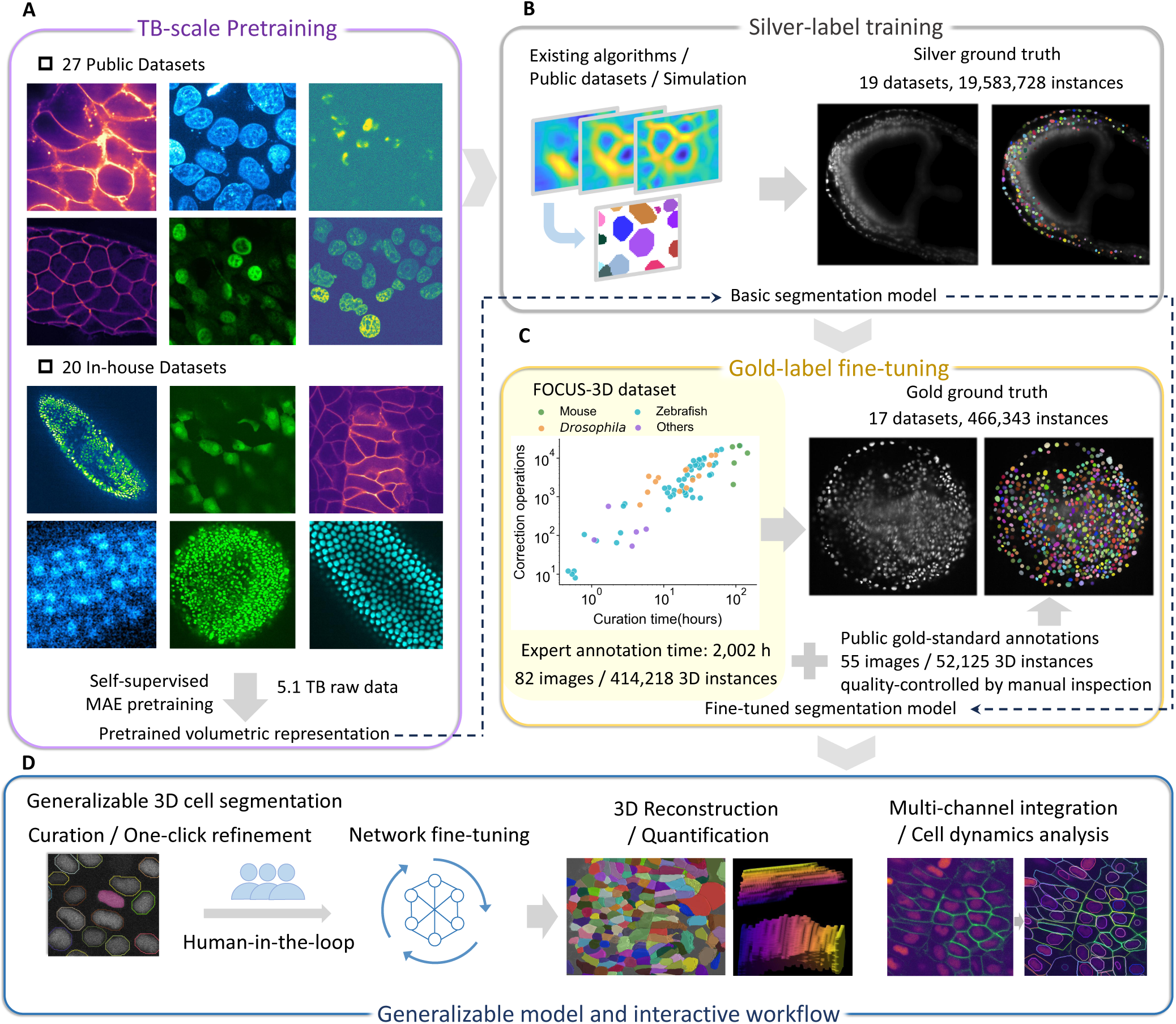
Datasets and training strategy for FOCUS-3D. (A) Self-supervised pretraining. This stage includes 27 public datasets and 20 in-house datasets, comprising a total of 5.1 TB of raw data. (B) Silver-standard training. Existing methods, simulation-based approaches and public datasets are used to generate 19,583,728 silver-standard annotations. (C) Gold-standard annotations for fine-tuning and evaluation. We constructed the FOCUS-3D dataset, consisting of 82 expert-curated volumes and 414,218 annotated 3D cell instances. This dataset was combined with 55 quality-controlled public gold-standard volumes containing 52,125 instances, resulting in a gold-standard annotation pool of 7 species, 137 volumes and 466,343 instances. (D) Interactive workflow and downstream applications. FOCUS-3D enables data visualization, segmentation curation, human-in-the-loop model fine-tuning, 3D reconstruction and quantification, cell tracking, multi-channel integration and cell dynamics analysis.

**Figure 2:**
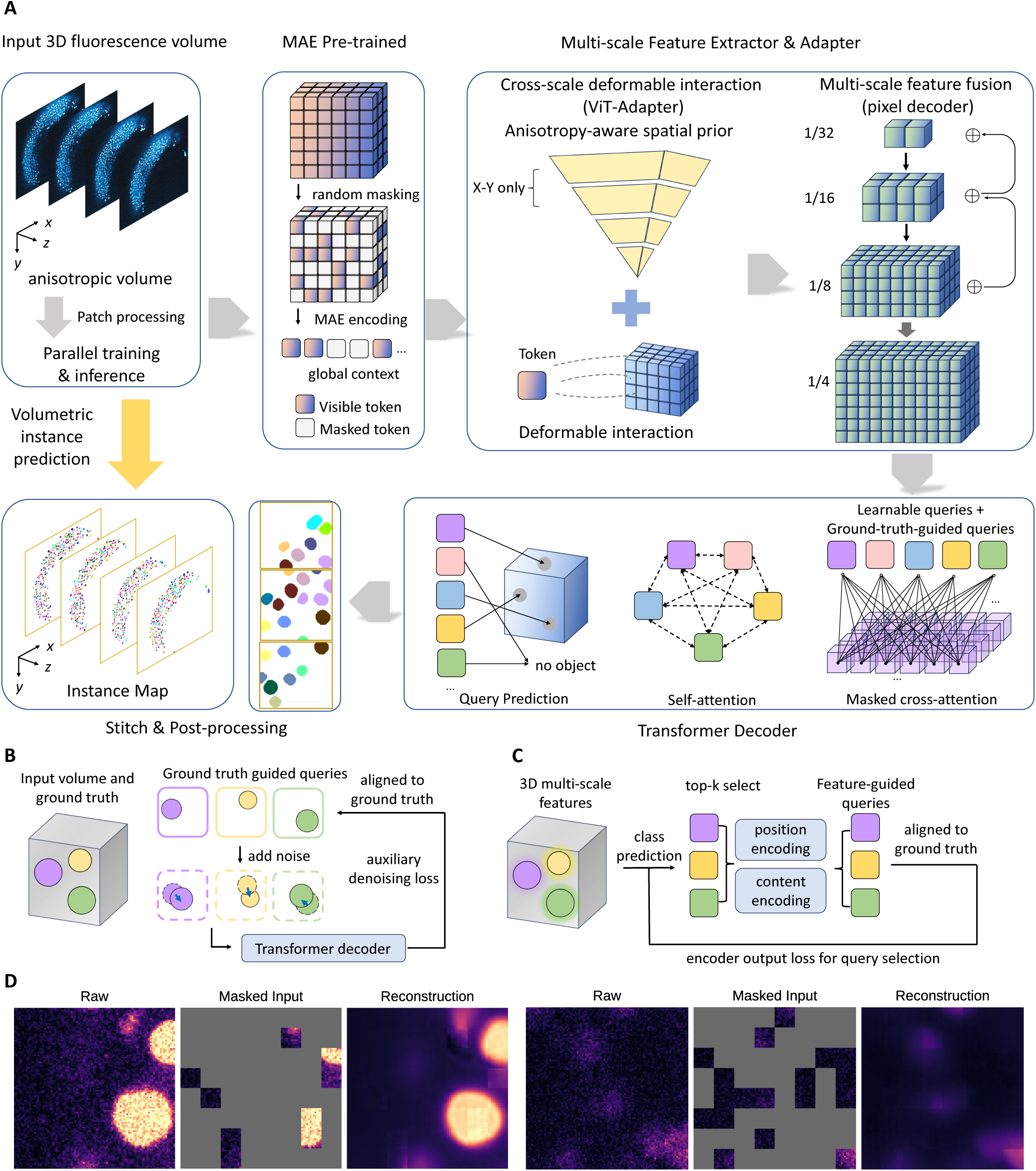
FOCUS-3D introduces a unified, integrated and end-to-end volumetric segmentation architecture for 3D cell segmentation. (A) Overall architecture of FOCUS-3D. The architecture consists of a 3D MAE backbone, a multi-scale feature extractor and adapter, a transformer decoder, and other key modules. (B) Schematic illustration of ground-truth-guided training. Auxiliary denoising loss is introduced to facilitate model training. (C) Schematic illustration of feature-guided query initialization. Queries are initialized by extracting the top-ranked candidates from multi-scale features. (D) Visualization of MAE reconstruction. Representative examples of the original image, masked input, and MAE reconst^8^ruction are shown.

We systematically validated FOCUS-3D across diverse species, tissues, fluorescent reporters, imaging modalities, and experimental setups. FOCUS-3D achieved state-of-the-art accuracy, generalizability, and computational efficiency, consistently outperforming existing methods on both in-domain and challenging out-of-domain testing data (Figure 3). The framework maintains robust performance despite substantial variations in image quality and experimental conditions. We further developed a user-friendly analytical workflow with FOCUS-3D integrated for visual inspection, correction, and dataset-specific refinement, enabling researchers to convert large microscopy volumes into curated cell-resolved datasets without requiring customized algorithm development^31,32^ (Figure 4). Released as a free, fully open-source platform, FOCUS-3D delivers both a powerful generalized segmentation model and a readily deployable analytical bridge that connects large-scale volumetric imaging to quantitative biological inquiry.

**Figure 3:**
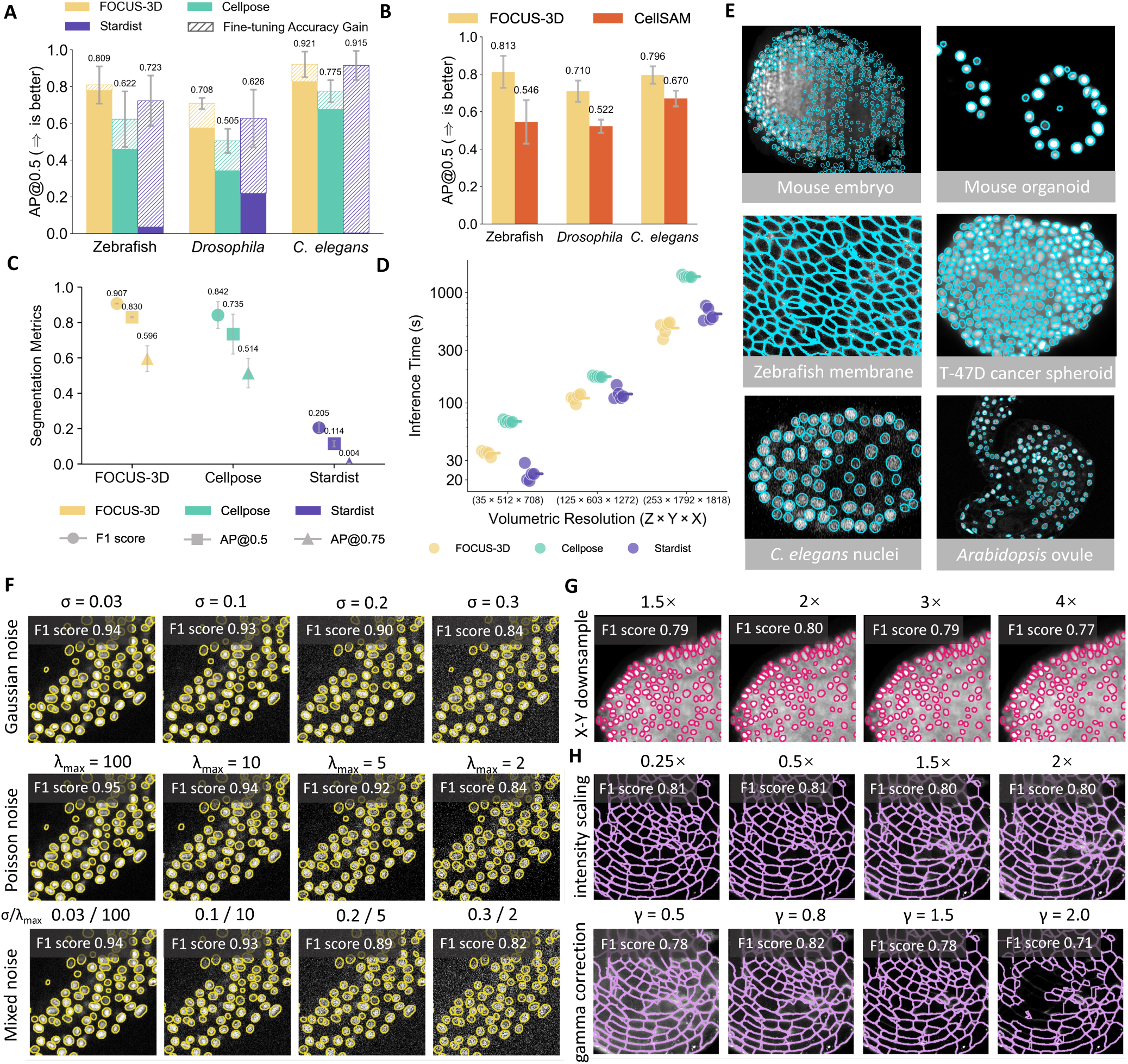
FOCUS-3D achieves state-of-the-art performance across diverse datasets. (A) Comparison of AP@0.5 before and after fine-tuning between FOCUS-3D and peer 3D cell segmentation algorithms. (B) AP@0.5 comparison between FOCUS-3D and the slice-by-slice 2D segmentation results of CellSAM. (C) Segmentation performance comparison among peer methods under the zero-shot setting on the *Arabidopsis* ovule dataset. (D) Comparison of segmentation inference time among peer methods under matched inference conditions. (E) Representative segmentation results of FOCUS-3D across diverse datasets. (F) Nuclear segmentation results of zebrafish embryos under different scales of Gaussian noise, Poisson noise, and mixed noise of Gaussian and Poisson noise. (G) Nuclear segmentation results of *Drosophila* embryos after upsampling following downsampling in the X-Y direction at different scales. (H) Membrane segmentation results of *Arabidopsis* ovule under linear transformation and gamma transformation of image intensity values at different scales.

**Figure 4:**
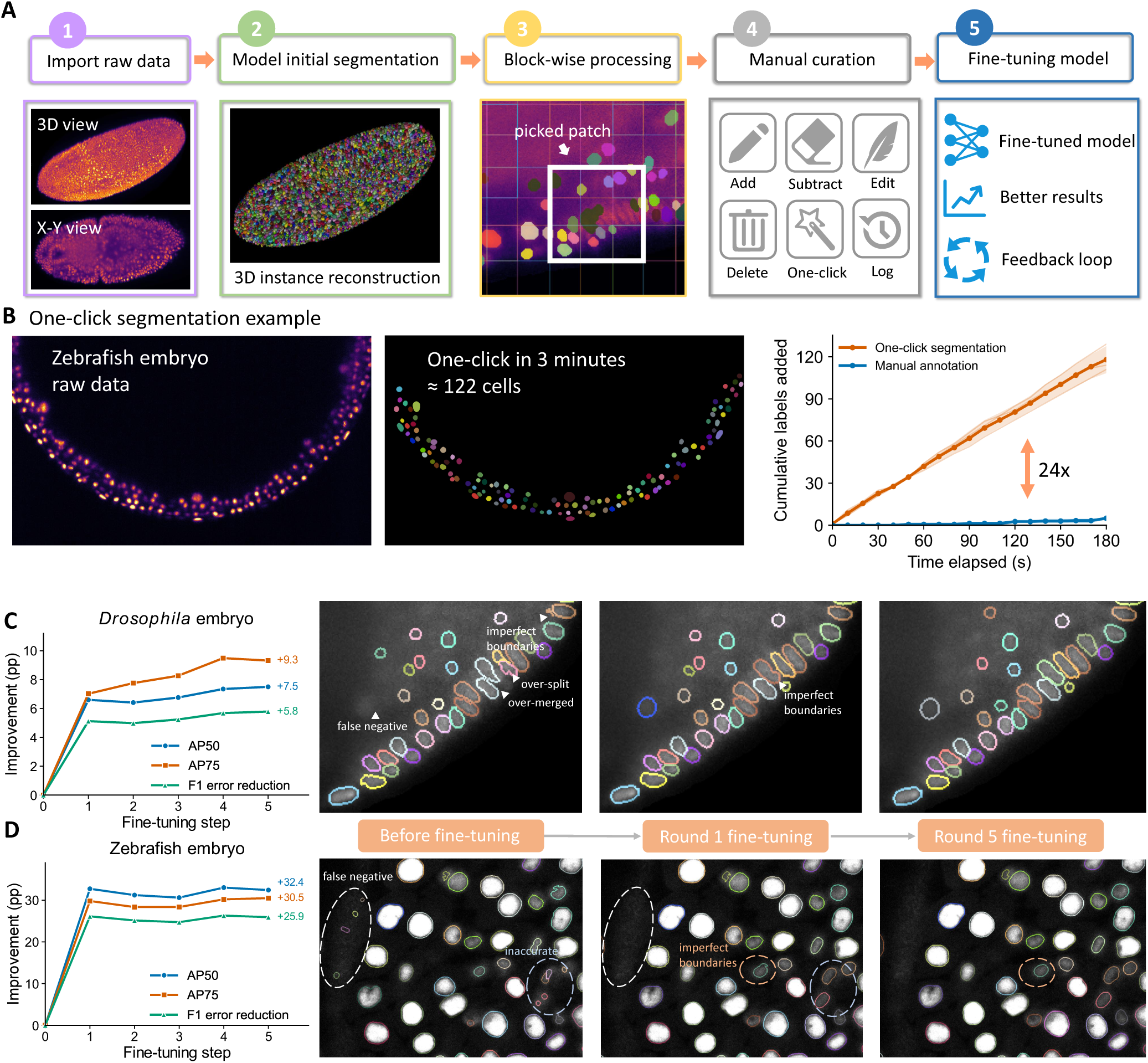
FOCUS-3D integrates efficient 3D cell curation with human-in-the-loop model adaptation. (A) The integrated workflow of FOCUS-3D, consisting of raw data import, model-based initial segmentation, block-wise processing, manual curation, and model fine-tuning. (B) Comparison of one-click segmentation and manual annotation on the zebrafish embryo dataset. One-click segmentation reduced the per-cell annotation time from 36 s to 1.5 s, corresponding to a 24-fold improvement in efficiency. (C) Human-in-the-loop results on *Drosophila* embryos. AP@0.5 and AP@0.75 improved by 7.5 and 9.3 percentage points, respectively, while the F1 error (1 − F1 score) decreased by 5.8 percentage points. (D) Human-in-the-loop results on zebrafish embryos. AP@0.5 and AP@0.75 improved by 32.4 and 30.5 percentage points, respectively, while the F1 error decreased by 25.9 percentage points.

To demonstrate FOCUS-3D’s capability for dynamic and integrative biological analysis, we examined dual-channel live imaging recordings of zebrafish notochord morphogenesis, in which nuclear signals encode cell movement and lineage information, while membrane fluorescence resolves cell morphology and tissue packing architecture^33^. By segmenting both the nuclear and membrane channels and associating nuclear tracking trajectories with membrane-defined cell boundaries^8^, we reconstructed cellular dynamic behaviors as well as comprehensive tissue-scale spatial organization. This high-resolution analysis uncovered three sequential developmental phases of notochord formation: compaction, consolidation, and deformation. We further discriminated early morphogenetic events driven by coordinated cellular motility from subsequent tissue remodeling stages governed by cell shape rearrangement and collective tissue repacking (Figure 5). We then used cell morphology as a shared phenotype to connect these longitudinal imaging states with spatial transcriptional states in a 6-somite zebrafish notochord^34^. Morphology-guided mapping identified distinct spatial counterparts of the early and late morphological states. The associated transcriptional differences were independently supported by developmental-stage-dependent single-cell RNA sequencing data^35^, linking dynamic 3D cellular remodeling to molecular state across independent datasets. These findings highlight that precise volumetric segmentation enables the discovery and quantitative characterization of subtle developmental processes inaccessible to manual visual inspection. By supporting robust, generalizable cellular analysis across heterogeneous, dynamic 3D microscopy datasets and enabling integration of cellular morphology with molecular state, FOCUS-3D provides a powerful foundational tool for dissecting how individual cellular behaviors collectively shape tissue architecture in development and pathogenesis.

**Figure 5:**
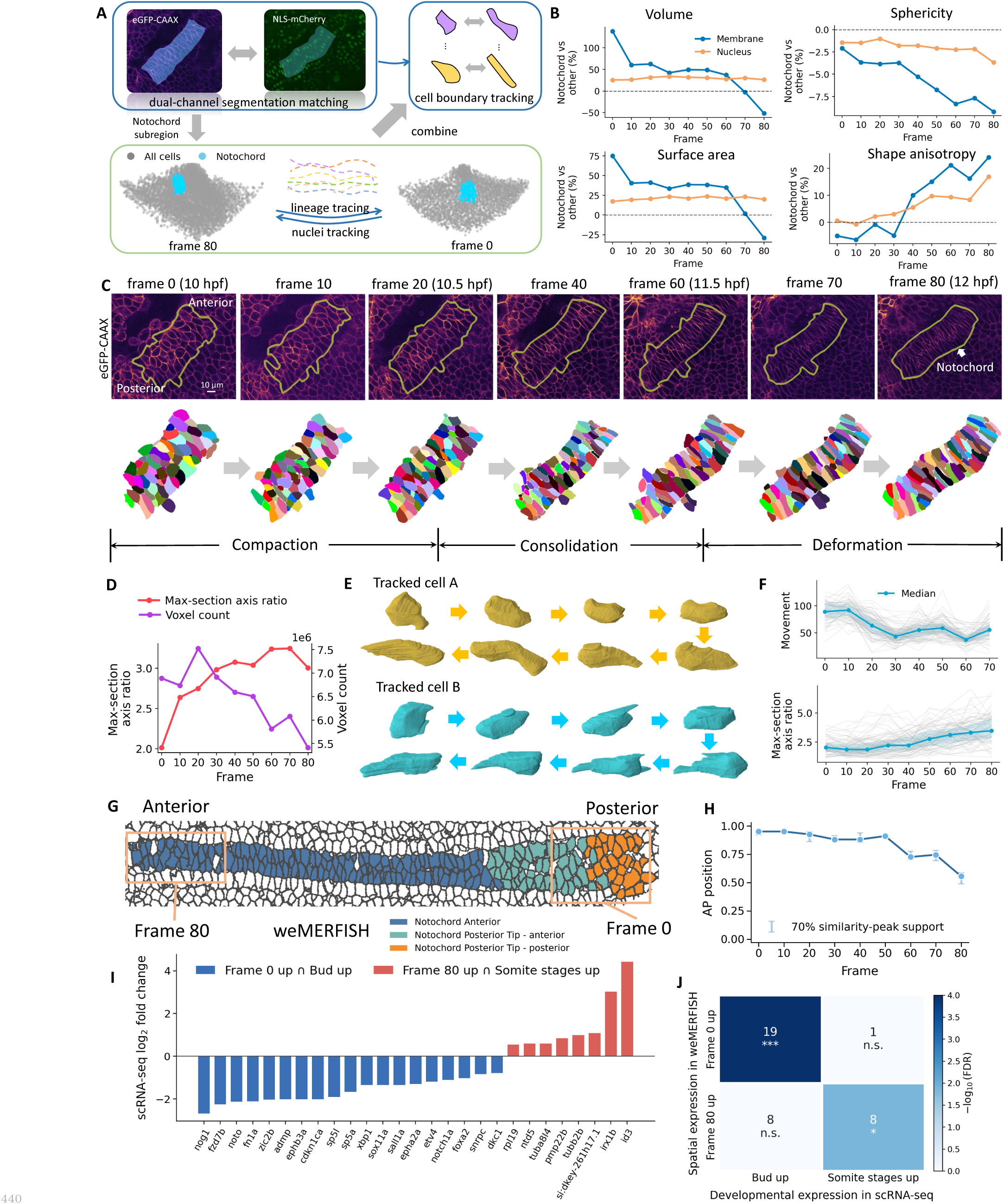
(A) Schematic of the dual-channel integration strategy used to convert nuclear trajectories into membrane-resolved cell-boundary trajectories. Nuclear and membrane channels were segmented independently with FOCUS-3D. Membrane instances were matched to nuclear positions at each time point, and ITEC-based^8^ nuclear tracking was then transferred to the corresponding membrane-defined cell boundaries. (B) Quantitative comparison of membrane- and nuclear-derived morphometric features between the lineage-defined notochord subregion and the surrounding cells in the same field of view. Relative differences in volume, surface area, sphericity, and shape anisotropy between notochord and surrounding cells were quantified separately from membrane and nuclear segmentations across time. (C) Temporal progression of the annotated notochord subregion from frame 0 (10 hpf) to frame 80 (12 hpf). Representative views show the evolution of the notochord region and the corresponding 3D cell segmentation, shown by colored cells. The same color across time denotes the same tracked cell. (D) Tissue-level morphometric dynamics of the tracked notochord subregion. The total segmented volume, calculated as the sum of anisotropy-corrected voxel volumes within the segmented cells, and the maximum-section axis ratio were measured over time. (E) Two representative examples of membrane-resolved single-cell dynamics. (F) Single-cell dynamic features within the notochord subregion after outlier exclusion. Median cell movement and median max-section axis ratio were plotted over time. (G) Spatial mapping of morphology-defined notochord states onto the 6-somite weMERFISH embryo. The notochord is shown together with the 10% AP regions providing the best morphology match to frame 0 and frame 80. (H) Morphology-guided temporal-to-spatial mapping along the anterior– posterior (AP) axis of the 6-somite weMERFISH notochord. Points indicate the center of the best-matching 10% AP window for each live-imaging frame. Error bars indicate the contiguous AP support interval surrounding the maximum morphology-similarity peak, defined using 70% of the peak prominence relative to the median similarity across the AP landscape. (I) Genes showing concordant spatial and temporal expression differences between the morphology-defined weMER-FISH regions and the independent single-cell RNA-seq analysis. Bars show the single-cell RNA-seq log_2_ fold change for the combined 3-somite/6-somite population relative to the bud stage; negative values therefore indicate higher expression at the bud stage and positive values higher expression at somite stages. Genes are grouped according to frame 0/bud-stage or frame 80/somite-stage concordance. (J) Directional overlap between morphology-defined weMERFISH spatial signatures and single-cell RNA-seq temporal signatures. Rows denote genes expressed more highly in the frame 0-like or frame 80-like weMERFISH regions, and columns denote genes upregulated at the bud stage or at somite stages in single-cell RNA-seq. Numbers indicate overlapping genes and color intensity represents *−* log_10_(FDR). Fisher’s exact tests used the 456 weMERFISH-measured genes matched and tested in the single-cell dataset as the gene universe, with Benjamini–Hochberg correction across the four directional comparisons.

## Results

### A large-scale, versatile and multi-tiered dataset is constructed for 3D cell instance segmentation

A central challenge in building an accurate and robust 3D cell instance segmentation model is the lack of large-scale, high-quality training datasets with broad coverage of diverse experimental conditions. Due to the extreme biological and technical heterogeneity, volumetric microscopy images vary substantially across organisms, tissues, cellular labels, imaging systems, spatial resolutions, and signal-to-noise conditions, whereas accurate 3D instance annotations remain scarce and costly to produce. Existing resources usually contain annotations for only a small number of cells in one or two specific scenarios. Assisted by our newly developed annotation platform, we overcame this challenge and constructed a large-scale dataset which consists of three tiers of data, unlabeled data for self-supervised representation learning, silver-standard machine-generated annotations for scalable supervised training, and expert-curated gold-standard annotations for high-quality model refinement and evaluation (Figure 1).

To capture the biological and technical diversity of 3D microscopy images, we first assembled arguably the most comprehensive dataset of unlabeled image volumes for self-supervised pretraining (Figure 1A). The pretraining dataset comprises approximately 5.1 TB of 3D microscopy data, including 47 sub-datasets and more than 17,000 image volumes collected from public repositories and in-house sources (Table S1). It covers diverse biological contexts across multiple species and tissue types, including plant tissues, yeast, *C. elegans*, *Drosophila*, zebrafish, mouse, human cells, and simulated cell populations. The dataset also spans major imaging categories, including point-scanning confocal microscopy, spinning-disk confocal microscopy, light-sheet microscopy, two-photon microscopy, and simulated fluorescence imaging. The images contain nuclear, membrane, and cytoplasmic signals acquired across different developmental stages, staining strategies, imaging resolutions, and signal-to-noise conditions. This collection provides broad coverage of volumetric structures and imaging conditions that would form a critical foundation to enable the learning of effective 3D cellular representation and assure the robustness across heterogeneous application scenarios.

At the intermediate annotation tier, we further constructed a silver-standard annotation dataset (Figure 1B). The goal of generating a silver-standard dataset is to leverage the efforts of the existing, though far-from-satisfactory, segmentation models. We first reused the machine-generated labels originally provided by the corresponding public sources. Otherwise, through a workflow combining unsupervised automatic segmentation and expert quality screening, we converted large amounts of unlabeled data into 3D instance annotations (see Table S2). For example, silver-standard annotations across 600 time points of zebrafish embryo development were generated using PrinCut-Auto^36^. Overall, this stage produced more than 19 million annotated instances across 19 datasets. By extending instance-level annotation far beyond the scale feasible for exhaustive manual curation, the silver-standard tier builds upon existing models and covers a broad range of cell morphologies, packing densities, tissue architectures, labeling targets, and image characteristics.

Finally, we established a gold-standard dataset through expert manual curation and stringent quality control, providing a high-fidelity reference for model development and evaluation (Figure 1C and Table S3). To our knowledge, it represents the largest gold-standard annotated dataset created specifically for 3D cell instance segmentation. Annotators manually inspected and corrected each 3D volume using our annotation platform, addressing errors such as missed cells, false-positive instances, boundary inaccuracies, and merged cells. To ensure annotation quality, the curated volumes then underwent an independent quality-control step, in which two additional reviewers randomly inspected the annotations, corrected remaining visible errors, and repeated this process until no obvious visually detectable errors were found. The entire annotation and correction process required over 2,000 person-hours and involved the annotation of more than 410,000 cell instances (Figure S1). For model development, we combined our curated dataset with quality-controlled public gold-standard annotations that passed manual inspection, resulting in a gold-standard training and evaluation pool of 137 image volumes and 466,343 annotated cell instances across 17 datasets.

Together, the three tiers form a large-scale and versatile dataset that spans both biological breadth and annotation fidelity: the unlabeled collection maximizes biological and technical coverage, the silver-standard tier scales instance-level annotation to millions of cells, and the gold-standard tier provides a rigorously curated reference for model refinement and evaluation. This multitiered resource provided the data foundation for the volumetric architecture and learning strategy described below.

### 3D unified, integrated, and end-to-end architecture was built for volumetric microscopy cell instance segmentation

Our model was built following three principles (Figure 2A). Firstly, volumetric cell segmentation should be performed natively in the three dimensions rather than reconstructed from independently segmented 2D views. Unified 3D modeling allows each cell to be interpreted using information distributed throughout its 3D neighborhood, which becomes indispensable when individual image planes are ambiguous because of dense packing, weak boundaries, or low signal-to-noise ratios. Moreover, slice-wise or orthogonal-view approaches require repeated inference over multiple image planes or views followed by additional cross-view reconciliation, introducing substantial computational overhead. Secondly, the framework should make effective use of data that are either unannotated or annotated at different levels of fidelity rather than relying exclusively on costly expert-curated 3D labels. While expert-curated 3D annotations are scarce and expensive, unlabeled volumetric images are abundant, and existing segmentation methods can provide imperfect but informative instance-level annotations. Therefore, we built an integrated architecture that could seamlessly utilize the widely available raw data for learning the essential 3D representation, the machine-generated imperfect labels for reusing the existing modeling efforts, and the scarce expert-curated cell labels for achieving the highest accuracy. Thirdly, as the segmentation rules are difficult to manually design, an end-to-end approach should be taken to learn the rules for directly predicting the cell instances from multi-scale spatial features. We draw on the query-based mask-classification idea introduced in Mask2Former^29^. Candidate cells are represented by object queries that directly produce instance-level volumetric masks. Because these queries decode from a shared multi-scale volumetric feature hierarchy, individual instances are predicted in the context of surrounding structures rather than in isolation. Features at different spatial scales further provide complementary cues for resolving cell-level structures under variations in cell size, morphology, and packing density.

In technical details, FOCUS-3D first uses the encoder of a 3D masked autoencoder (MAE) as its volumetric backbone^37^, which was modified from the original 2D version. The backbone converts input volumes into context-rich token representations that preserve information across the three spatial dimensions (Figure 2D). The MAE, conceptually similar to self-supervised representation learning in large language models^38–40^, is to take advantage of the widely available raw volumetric images and learn the basic image representations to enhance both segmentation accuracy and robustness. Then ViT-Adapter converts the single-scale MAE representations into hierarchical multi-scale feature maps^41^. To accommodate the anisotropic resolution of 3D microscopy images, FOCUS-3D employs an anisotropy-aware downsampling strategy that performs downsampling only in the X-Y dimensions, while keeping the Z dimension fully resolved throughout the early feature pyramid stages. The 3D pixel decoder consolidates these hierarchical representations into high-resolution mask features^29^, progressively increasing spatial discrimination and instance separability before final decoding (Figure S2). The transformer decoder then treats segmentation as object-level set prediction, with each query producing a cell score and a complete 3D mask^29^. The weights in the decoder were initialized through training on the machine-generated labels and then refined on the expert-curated annotations with a smaller learning rate. Importantly, instance representations and mask predictions are learned jointly from the volumetric feature hierarchy, making object formation data-driven rather than dependent on predefined cell models or hand-crafted segmentation rules. For full-volume inference, overlapping volumetric blocks are processed through the same pathway and stitched into a global instance map. Together, these components establish a continuous volumetric prediction pathway from 3D feature representation to complete cell-instance reconstruction.

### Architectural innovations improve model accuracy

Although the unified architecture establishes a feasible end-to-end volumetric prediction pathway, directly applying the conventional query-based decoding to dense 3D cell images is challenging. Input-agnostic queries provide limited information about where candidate cells are located, whereas the large number of similar and closely packed instances makes query assignment and decoder optimization difficult. We therefore introduced two complementary strategies targeting query initialization and decoder training, respectively. Instead of relying solely on input-agnostic learned queries, FOCUS-3D initializes object queries from top-ranked encoded volumetric features^30,42,43^ (Figure 2C), allowing the initial queries to carry image-dependent content and spatial priors. This procedure anchors the initial queries to image-dependent candidate regions before iterative decoding, providing the decoder with more informative starting points for instance prediction. In addition, during training, FOCUS-3D introduces ground-truth-guided queries with controlled perturbations as an auxiliary denoising task (Figure 2B). By learning to recover the corresponding instances from perturbed queries, the decoder can be better trained as it receives more direct supervision for query assignment and iterative mask prediction.

To validate the contribution of these design choices, we performed two ablation experiments. Model performance was quantified using three widely used metrics, AP@0.5 (average precision at a 50% intersection-over-union threshold), AP@0.75 (average precision at a 75% intersection-over-union threshold) and F1 score. To provide a stringent test of these architectural choices, we evaluated all model variants in a cross-dataset setting, where models trained on a mixed training set consisting of zebrafish, *Drosophila*, and *C. elegans* embryo datasets were tested on zebrafish embryo data acquired under a different imaging condition (Figure S3). Removing feature-guided query initialization caused a marked performance drop, with AP@0.5 decreasing from 0.583 to 0.428. Removing denoising queries also reduced AP@0.5 from 0.583 to 0.564, with a more pronounced decrease under the stricter AP@0.75 criterion, from 0.409 to 0.385. Together, these ablations confirm that both feature-guided query initialization and ground-truth-guided training contribute to segmentation accuracy.

### Multistage training improves accuracy and generalization with limited annotation

Volumetric microscopy data are abundant, but expert-curated 3D instance annotations remain practically limited. We therefore designed a multistage training strategy that uses large-scale unlabeled and silver-standard data to amplify the value of the scarce gold-standard supervision. Self-supervised masked autoencoder (MAE) pretraining^37^ first captured the shared volumetric structures from the unlabeled microscopy data without requiring any instance annotations. The pretrained model was then trained on the silver-standard annotations to learn instance-level prediction across a broad range of biological structures and imaging conditions. Finally, the gold-standard annotations were used to refine the model against rigorously curated 3D cell instances. This multistage design integrates general representation learning, scalable instance-level learning, and high-fidelity model refinement, allowing abundant but imperfect data to complement rather than replace expert-curated annotations.

We first evaluated the contribution of MAE pretraining under different annotation budgets using 20%, 50%, or 100% of a mixed gold-standard training set comprising zebrafish, *Drosophila*, and *C. elegans* embryo datasets. Models initialized from the MAE-pretrained backbone consistently outperformed models with randomly initialized backbone parameters. The impact of MAE was greater when the model was applied to out-of-domain datasets that were not in the gold-standard training set (Figure S4A), where the improvement of AP@0.5 was 0.01 for the in-domain applications, compared to an improvement of 0.129 for the out-of-domain applications, from 0.454 with random initialization to 0.583 with MAE pretraining. Furthermore, the improvement was greatest when annotated training data were limited: MAE pretraining increased AP@0.5 by 0.224 under the 20% training-data setting, compared with gains of 0.202 and 0.129 when 50% and 100% of the training data were used, respectively. Therefore, large-scale self-supervised pretraining is generally useful and particularly evident when the availability of expert annotations is limited or the model is generalized to different settings of experiments.

Next, we examined the impact of learning from the silver-standard datasets. Using the same evaluation setting, we compared three training configurations starting from the same MAE-pretrained backbone: gold-standard training alone, silver-standard training alone, and silver-standard training followed by gold-standard refinement (Figure S4B). Training directly on the gold-standard data achieved an AP@0.5 of 0.583, whereas introducing large-scale silver-standard training before gold-standard refinement increased AP@0.5 to 0.785, an absolute gain of 0.202. Notably, despite receiving no gold-standard supervision, MAE pretraining followed by silver-standard training alone achieved an AP@0.5 of 0.748 on the unseen evaluation data, emphasizing the value of broadly leveraging existing modeling efforts in the field. Subsequent gold-standard refinement further increased AP@0.5 from 0.748 to 0.785, confirming the additional value of high-fidelity expert annotations. These results show that silver-standard supervision provides substantial transferable instance-level information across datasets, while gold-standard annotations further refine this broadly learned representation toward higher segmentation accuracy.

### FOCUS-3D outperforms peer methods in both accuracy and robustness across diverse and unseen biological scenarios

To evaluate the overall performance across diverse volumetric microscopy datasets, we benchmarked FOCUS-3D against two widely used peer methods, Cellpose^44^ and StarDist^26^, under both generalist and gold-standard fine-tuned settings using three datasets with distinct cellular morphologies and imaging characteristics: zebrafish, *Drosophila*, and *C. elegans*.

We first evaluated generalist models without target-dataset fine-tuning. For each benchmark dataset, performance was reported under a five-fold evaluation protocol, with a subset of 3D volumes held out for testing in each fold. FOCUS-3D used weights trained with large-scale silver-standard annotations but excluded gold-standard annotations from the target benchmark dataset, whereas Cellpose and StarDist used their publicly released pretrained models. In this setting, FOCUS-3D achieved the best overall segmentation performance across the three benchmarks (Figure 3A and Figure S5A). On the *Drosophila* benchmark, FOCUS-3D achieved a mean AP@0.5 of 0.575, outperforming Cellpose (AP@0.5 = 0.342) and StarDist (AP@0.5 = 0.218). After fine-tuning with gold-standard annotations from the corresponding training split, FOCUS-3D further improved its performance and achieved a mean AP@0.5 of 0.708 on the *Drosophila* benchmark, exceeding Cellpose and StarDist by 0.203 and 0.082, respectively. Together, these results demonstrate that FOCUS-3D provides accurate volumetric segmentation across diverse biological systems.

We then compared FOCUS-3D with CellSAM^16^, a recently developed foundation model for 2D cellular image segmentation (Figure 3B and Figure S5B). To enable a direct comparison, we evaluated 2D cross-sections extracted from FOCUS-3D predictions against CellSAM’s native slice-level outputs. Across the zebrafish, *Drosophila* and *C. elegans* datasets, FOCUS-3D achieved AP@0.5 scores of 0.813, 0.710 and 0.796, respectively, whereas CellSAM achieved AP@0.5 scores of 0.546, 0.522 and 0.670. These results indicate that the segmentation accuracy achieved by FOCUS-3D remains evident even when evaluated on individual image planes, suggesting that direct volumetric modeling can better preserve fine-scale object boundaries and structural information than using the individual slices independently.

Representative examples illustrate that FOCUS-3D can delineate diverse biological structures, including cancer spheroids, zebrafish membranes, and mouse organoids (Figure 3E). We next assessed the ability of FOCUS-3D to generalize beyond the biological systems represented in the training data. To this end, we evaluated the model under a zero-shot setting using an independent *Arabidopsis* ovule dataset^45^ that was not used during either pretraining or fine-tuning (Figure 3C). Despite differences in tissue architecture, FOCUS-3D maintained consistently high segmentation accuracy across all tested subsets, achieving a mean AP@0.5 of 0.830 across the three images. In comparison, Cellpose achieved a mean AP@0.5 of 0.735, whereas StarDist achieved a mean AP@0.5 of 0.114. Although Cellpose achieved competitive performance on individual subsets, its performance varied considerably among individual test images even within the same dataset, as shown by the gray error bars in Figure 3C. In contrast, FOCUS-3D maintained stable and accurate predictions without task-specific retraining or parameter optimization, demonstrating strong cross-domain generalization.

### Fast, robust volumetric segmentation empowers practical, everyday deployment in real-world experimental workflows

Practical volumetric segmentation requires both computational efficiency and stable performance under variable imaging conditions. We first benchmarked inference efficiency under a standardized 12-GB VRAM budget (Figure 3D). By operating directly on volumetric inputs, FOCUS-3D avoids slice-by-slice processing and enables efficient full-volume segmentation, completing inference substantially faster than Cellpose. On a zebrafish dataset, for example, FOCUS-3D completed inference in 477.7 s, compared with 1402.3 s for Cellpose, corresponding to approximately 3-fold faster inference. These results show that accurate full-volume segmentation can be performed efficiently within a resource-limited workstation setting.

Beyond inference efficiency, we assessed the stability of FOCUS-3D under controlled degradations that mimic common variations in microscopy image quality. To test sensitivity to image noise, we progressively introduced synthetic Gaussian, Poisson, and mixed noise to the zebrafish embryo dataset (Figure 3F). FOCUS-3D retained stable instance-level performance across the tested noise levels, with an F1 score of 0.82 under the strongest mixed-noise condition tested (*σ* = 0.3, *λ_max_* = 2). We further assessed sensitivity to lateral resolution degradation by down-sampling *Drosophila* embryo volumes in the X-Y plane (Figure 3G). Across scaling factors up to 4*×*, FOCUS-3D largely preserved the overall morphology of predicted masks, with F1 scores remaining between 0.77 and 0.80. To assess the effect of intensity variation, we applied linear intensity scaling and non-linear gamma transformations to *Arabidopsis* ovule membrane images (Figure 3H). Although performance decreased to an F1 score of 0.71 under an extreme gamma transformation, the model retained excellent segmentation performance under the less extreme intensity perturbations. Together, efficient full-volume inference and resilience to noise, resolution loss, and intensity shifts show that FOCUS-3D is not confined to ideal benchmark conditions. It maintains stable segmentation across the computational and image-quality variations that must be accommodated in the real-world volumetric microscopy experimental conditions.

### One-click correction accelerates 3D cell segmentation inspection and curation within an integrated platform

Although general 3D segmentation models can provide reliable initial results across diverse microscopy datasets, biological images under special experimental settings can exhibit substantial variations in tissue morphology, imaging noise, fluorescence intensity distributions, and cell density. As a result, fully automated segmentation pipelines may need to have the results inspected and corrected when applied to these scenarios. To facilitate efficient result inspection and correction, we developed an integrated interactive platform for 3D cell segmentation and curation.

Implemented as a napari plugin^46^, the platform consists of raw 3D image loading, model-based segmentation, block-wise curation, model fine-tuning, re-segmentation, and quantitative analysis (Figure 4A and Figure S6). Biologists first apply a publicly available pretrained model to obtain an initial full-volume segmentation. They can then select local patches in regions of interest or regions with frequent errors for inspection and correction. The plugin supports a series of commonly used 3D curation operations, including adding/subtracting voxels, creating new instance labels, deleting erroneous instances, and batch deletion within a user-defined region. The curated labels can be reused for dataset-specific model fine-tuning, as described in the next section.

To reduce the cost of local curation, we designed a one-click correction function. For cells that are missed, incomplete, or incorrectly segmented by the automatic model, users only need to click a point inside the target cell, after which the model automatically generates the corresponding 3D instance segmentation based on the spatial prompt (see Supplementary Notes). To quantitatively evaluate the efficiency of cell-wise curation, we benchmarked the time required to complete individual 3D cell instances using one-click correction or conventional manual annotation, with each approach independently repeated three times (Figure 4B). Compared with conventional slice-by-slice manual editing, one-click correction substantially reduces the number of user interactions and better matches the practical requirements of correcting cell instances in 3D images. On the zebrafish embryo dataset, the average time required to complete a volumetric cell instance was approximately 1.5 s using one-click correction, compared with approximately 36 s using conventional manual annotation. One-click correction reduced the per-cell curation time by approximately 24-fold. Thus, the integrated platform combines flexible manual editing with rapid one-click correction, substantially reducing the effort required to curate volumetric cell segmentations.

### FOCUS-3D is adaptable to new datasets with minimal curation through human-in-the-loop refinement

Beyond correcting individual segmentation errors, the curated labels generated within the platform can be fed back into FOCUS-3D for dataset-specific model refinement. Users can fine-tune the pretrained model on selected curated blocks and apply the updated model to re-segment the complete volume, forming a closed loop between prediction, expert curation, and model adaptation^31^.

To determine whether this closed-loop refinement could adapt the model with limited curation, we evaluated the human-in-the-loop fine-tuning process on two representative 3D microscopy datasets (Figure 4C and Figure 4D). Starting from the initial segmentation generated by the pretrained model, we performed five rounds of fine-tuning. In each round, expert-curated training blocks were randomly selected to cover 2% of the entire image and were then added to update the model, followed by re-evaluation of segmentation performance on the full image. In the zero-shot evaluation on *Drosophila* embryos, fine-tuning increased AP@0.5 and AP@0.75 by 7.5 and 9.3 percentage points, respectively, while reducing the F1 error (1 − F1 score) by 5.8 percentage points. After refinement, densely packed cells under low signal-to-noise conditions were more accurately separated. In another zebrafish embryo dataset, where cells exhibited substantial variations in intensity distributions, AP@0.5 and AP@0.75 improved by 32.4 and 30.5 percentage points, respectively. This indicates that even when the initial model fails to accurately segment cells with relatively low intensities, fine-tuning provides sufficient adaptability to the new data distribution. Notably, in both examples, only 2% of the data was sufficient to produce a substantial performance gain, greatly reducing the curation burden for users.

These experiments show that human-in-the-loop refinement can leverage expert corrections from only a small fraction of the data to improve segmentation across the complete volume. By feeding curated labels back into the model, this strategy adapts the pretrained model to new settings with dataset-specific morphology, signal quality, and intensity distributions. More broadly, the closed loop linking initial prediction, expert curation, model updating, and re-segmentation converts localized expert input into dataset-wide model adaptation, providing a practical path from general-purpose segmentation to accurate dataset-specific results.

### FOCUS-3D reconstructs cell-membrane dynamics by integrating with nuclear tracking

A central challenge in developmental morphodynamics is to confidently follow the same cells while their 3D boundaries, positions and neighborhood relationships are continuously remodeled^33,47^. Nuclear tracking provides robust cell identity and motion trajectories^8,48^, but it does not provide information about membrane-defined geometry that determines cell shape and packing. Conversely, membrane segmentation captures the physical cell boundary, but without a stable lineage coordinate it is difficult to interpret shape changes of the same cells over long volumetric time series^33^. Although FOCUS-3D is designed as a general 3D cell segmentation framework, its ability to segment both nuclear and membrane channels provides the foundation for linking these complementary representations. By combining FOCUS-3D membrane and nuclear segmentation with nuclear tracking, we converted nuclear trajectories into membrane-resolved cell-boundary trajectories through nucleus-to-membrane matching in the same cell. This can enable, for instance, analysis of organ morphogenesis during complex processes such as gastrulation and subsequent stages that otherwise would be difficult or burdensome.

We used zebrafish notochord morphogenesis as a representative application of the capabilities of FOCUS-3D. The notochord is a classical model of axial tissue remodeling, in which a broad cellular field is progressively reorganized into a narrow, elongated midline structure through convergence-extension-associated cell rearrangement^47,49–51^. This process cannot be solely described from nuclear positions alone: tissue narrowing, cell intercalation, local multilayer-to-bilayer transitions and membrane deformation all require cell-boundary-resolved measurements in three dimensions across time. We therefore applied FOCUS-3D to an 81-frame dual-channel time series spanning approximately 10–12 hpf (from the bud to the 6-somite stage), segmented both nuclei and membranes, and used ITEC-based nuclear tracking^8^ as the cell lineage scaffold. Each tracked nucleus was then assigned to its enclosing membrane instance, yielding longitudinal trajectories of cell volume, surface area, shape anisotropy, maximum-section axis ratio, cell displacement and local tissue organization (Figure 5A). To define a biologically coherent notochord population at every frame, we annotated a compact notochord subregion at frame 80 (12 hpf) and traced its nuclear ancestor cells backward through the time series. The recovered ancestor cells originated from a spatially defined cluster at 10 hpf, indicating that the late-stage annotation corresponded to a coherent notochord progenitor cell population rather than a transient geometric selection. Relative to the remaining cells in the same imaging field, the lineage-defined notochord population showed dynamic changes in morphological features over time (Figure 5B and Figure S7). At the population level, notochord cells became progressively smaller and less spherical than other adjacent cells, as reflected by decreasing relative cell volume, surface area, and sphericity, while their shape anisotropy increased (see Supplementary Notes for metric definitions). In contrast, nuclear measurements showed weaker relative differences overall, including a less pronounced increase in shape anisotropy. This integration therefore extends nuclear trajectories from positional measurements to membrane-resolved descriptions of cell shape and tissue organization over time.

### Lineage-resolved membrane dynamics reveal three phases of zebrafish notochord remodeling

Using the lineage-resolved membrane representation, we next examined the temporal evolution of this lineage-defined notochord population in greater detail. The lineage-defined notochord population followed a distinct membrane morphodynamic trajectory. At early time points, these cells had larger membrane volume and surface area than non-notochordal neighboring regions, consistent with a relatively loose and less compact local organization. Two transitions then became apparent: the first occurred between frame 0 and frame 20, when the notochord region narrowed and extended while individual cell volume remained comparatively stable, suggesting that early compaction was dominated by cell displacement and neighbor rearrangement rather than by simple cell shrinkage. Between frame 20 and frame 60, cell displacement slowed and membrane-derived features changed more gradually, consistent with progressive consolidation of the local cellular arrangement. A second transition emerged around frame 60 (11.5 hpf), after which membrane volume and surface area decreased more sharply and became lower than those of surrounding cells, indicating a late shift toward tighter packing and stronger cell-boundary remodeling (Figure 5B).

These two transition points partitioned the remodeling trajectory into three morphodynamic phases (Figure 5C). In the compaction phase, from approximately 10 to 10.5 hpf, cells showed pronounced displacement while maintaining relatively stable volume, and the tissue field narrowed along one axis while extending along the axial direction. This supports a movement-dominated compaction mode in which collective rearrangement precedes major membrane deformation. In the consolidation phase, from approximately 10.5 to 11.5 hpf, cell motion slowed, the population became more regularly arranged, and membrane features changed gradually, consistent with progressive alignment and stabilization of local packing. In the deformation phase, from approximately 11.5 to 12 hpf, the strongest cell shape remodeling occurred (Figure 5E). Cells shifted from a more multilayered arrangement toward a predominantly bilayered organization, with local regions retaining three or more layers, and this transition coincided with sharper changes in membrane surface area, maximum-section axis ratio and displacement (Figure 5B, Figure 5F and Figure S7).

Together, these measurements resolve notochord remodeling as a temporally ordered process in which movement-dominated compaction is followed by consolidation and later cell deformation and repacking. More broadly, integrating stable nuclear identities with membrane-resolved cell geometry provides a unified framework for connecting cell trajectories, shape remodeling, and tissue-scale reorganization across long volumetric time series. This integration bridges the complementary strengths of nuclear tracking and membrane segmentation, allowing cellular identity and motion to be analyzed together with the evolving physical geometry of the same cells. It therefore provides a general strategy for transforming multichannel 3D time-lapse imaging into quantitative, lineage-aware descriptions of cellular and tissue morphodynamics.

### Morphology-guided integration links notochord remodeling to transcriptional state change

The pronounced remodeling of notochord cell shape raised the possibility that the morphological states resolved by live imaging might also reflect changes in cellular state. To explore this connection, we used cell morphology as a bridge between the time-resolved imaging data and a 6-somite stage spatial transcriptomics (weMERFISH) dataset^34^. We first represented cells in both datasets using a common set of morphological features describing cell size and shape, and compared their feature distributions using an equally weighted Wasserstein-distance-based similarity score. By scanning local windows along the anterior–posterior (AP) axis of the spatial dataset, we identified the region that most closely matched the morphology observed at each time point. Strikingly, the morphology-defined frame 0 state mapped to a region containing only posterior-tip populations, whereas the frame 80 state mapped to a region containing only the anterior population, consistent with the existing weMERFISH notochord annotations (Figure 5G, Figure 5H and Figure S8). Thus, the temporal morphological transition observed within the same tracked lineage had distinct spatial analogues within the somite-stage tissue.

We next asked whether these morphology-defined regions were also associated with distinct molecular states. Comparison of their weMERFISH measured gene expression profiles revealed broad transcriptional differences between the frame 0-like and frame 80-like regions (Figure S9A). The frame 0-like region showed higher expression of genes associated with notochord identity and early tissue organization, including *noto*, *foxa2*, the developmental signaling gene *notch1a*, the extracellular matrix gene *fn1a* and the Wnt-responsive transcription factor *sp5l*. In contrast, the frame 80-like region showed higher expression of genes associated with later cell-state regulation and intracellular organization, including *id3*, *irx1b* and the microtubule gene *tubb2b* (Figure S9).

Because these spatial transcriptional differences were identified within a single 6-somite embryo, we next directly tested whether they reflected developmental progression using an independent single-cell RNA-seq dataset spanning the bud, 3-somite and 6-somite stages^35^. Gene Ontology analysis of the genome-wide stage-dependent expression changes showed that genes expressed more highly at the bud stage were enriched for RNA processing and splicing, chromatin remodeling and Wnt signaling, whereas genes expressed more highly at the 3-somite/6-somite stages were enriched for extracellular matrix and collagen organization, notochord development and notochord morphogenesis (Figure S10). Genes enriched in the weMERFISH frame 0-like region were significantly enriched among genes expressed more highly at the bud stage in the notochord (one-sided Fisher’s exact test, odds ratio = 5.06, *P* = 1.1 *×* 10*^−^*^5^, FDR = 4.2 *×* 10*^−^*^5^), whereas genes enriched in the weMERFISH frame 80-like region were significantly enriched among genes upregulated at somite stages (odds ratio = 7.01, *P* = 6.97 *×* 10*^−^*^3^, FDR = 1.39 *×* 10*^−^*^2^) (Figure 5J). Neither reverse comparison was significant. Thus, independent temporal transcriptomic data supported the developmental direction inferred from morphology-guided spatial mapping, linking the early and late morphological states to corresponding temporal transcriptional programs.

Together, these results connect the membrane-resolved morphodynamic trajectory to transcriptional state across independent datasets. The early and late morphological states identified from longitudinal imaging have distinct spatial counterparts in the 6-somite notochord, and the direction of the transcriptional differences between these regions is independently supported by developmental-stage-dependent expression changes in single-cell RNA sequencing. Rather than treating AP position as a direct proxy for developmental time, this analysis uses morphology as an intermediate phenotype to connect dynamic 3D cell behavior with molecular state, illustrating how FOCUS-3D can extend longitudinal imaging beyond morphodynamics toward integrative analysis of cellular phenotype and gene expression.

## Discussion

3D cell segmentation for volumetric microscopy data has long been a challenging problem in bioimage analysis^3–5^. Owing to the high cost and difficulty of 3D annotation, the field has lacked large-scale annotated datasets for many years^23,25^. To address this limitation, we first constructed a large voxel-level gold-standard dataset comprising 82 volumetric images and more than 410,000 annotated cell instances, and further developed a three-stage training strategy to improve model generalization across diverse datasets. These efforts ultimately led to the development of a generalist model for 3D cellular imaging data. We believe that the release of FOCUS-3D and its accompanying large-scale dataset will substantially promote progress in this field, facilitate community exchange, and drive algorithmic development. We also found that large-scale silver-standard annotations, when sufficiently diverse and quality-controlled, can provide useful segmentation priors and support strong performance even before gold-standard fine-tuning.

During the development process of this foundational model, we recognized that large-scale pre-trained volumetric representations are important for improving generalization across heterogeneous microscopy data. Methodologically, we extended and adapted segmentation methods developed for natural images according to the characteristics of 3D biological images^29^, and proposed a dedicated architecture for 3D cell segmentation, achieving leading performance. In particular, the MAE-based ViT-large backbone captures broadly shared volumetric features^11,37^, whereas multi-scale feature fusion and query-based decoding adapt these representations to image-specific cell morphology and boundary structure.

Building on a model with high accuracy and strong generalization, we further developed an integrated platform to accommodate diverse user-specific needs for 3D segmentation and improve the practical usability of the overall framework. We expect that FOCUS-3D will allow users to complete the full workflow, from image preprocessing and model fine-tuning to complete segmentation, thereby forming a closed-loop analysis pipeline.

Accurate 3D cell segmentation underpins large-scale cell tracking and lineage reconstruction^6–8^. Integrating nuclear and membrane channels, FOCUS-3D enables membrane-resolved morphodynamic analysis beyond static segmentation. Our analysis reveals that zebrafish notochord morphogenesis proceeds through three distinct phases—compaction, consolidation, and deformation— each defined by a unique interplay between cell motility and cell shape remodeling. Compaction is driven primarily by collective cell rearrangement, followed by consolidation during which cells progressively refine neighbor contacts. The final deformation phase exhibits the most pronounced membrane remodeling. These results demonstrate that integrating directly measured cellular behaviors can reveal temporally distinct phases of tissue morphogenesis. These phenotypic transitions were further linked to molecular state by using cell morphology as an intermediate phenotype to integrate the longitudinal imaging data with spatial and temporal transcriptomic datasets^34,35^. Early and late morphological states mapped to distinct regions of the 6-somite notochord with different transcriptional signatures, and the direction of these differences was independently supported by developmental-stage-dependent expression changes in single-cell RNA sequencing. In the future, more accurate long-term tracking, improved detection of cell division events, analysis over larger fields of view, and integration with temporally matched molecular measurements will further reveal how cell movement, division, shape change, neighbor rearrangement, and changes in cellular state collectively drive tissue morphogenesis. Together with emerging approaches that connect developmental cell behaviors with molecular identities^52–56^, our results illustrate how quantitative 3D morphology can serve as an intermediate phenotype for bridging longitudinal imaging and transcriptomic measurements. These efforts will become of vital importance in other events with defined phases, such as tissue regeneration^57–59^.

Several limitations remain. Although FOCUS-3D generalizes across a range of tested microscopy datasets, performance may decrease when new data differ substantially from the training distribution in imaging modality, staining pattern, cell size, or signal-to-noise ratio. Dataset-specific fine-tuning with a small number of curated annotations may therefore still be required. Segmentation errors can also occur under weak or discontinuous boundaries, strong intensity heterogeneity, densely touching cells, and extreme object sizes, where small cells may be missed or merged and large or elongated cells may be fragmented. In addition, the current framework is primarily designed and evaluated for nuclei and membrane-defined cell instances. Extending it to other biological structures, such as vesicles, organelles, or subcellular condensates, will require task-specific annotations, scale-aware adaptation, and dedicated evaluation. Finally, downstream biological analyses, including lineage-aware membrane morphodynamics, depend not only on segmentation quality but also on tracking accuracy, nucleus-to-membrane matching, and manual quality control. The morphology-guided transcriptomic integration also has several limitations. The spatial transcriptomic comparison was performed within a single 6-somite embryo, and the morphology-matched spatial regions should therefore be interpreted as spatial analogues of the longitudinal imaging states rather than as direct temporal equivalents. Moreover, the live-imaging, spatial transcriptomic, and single-cell RNA-seq measurements were obtained from independent datasets rather than from matched cells or regions in the notochord. Although the independent single-cell RNA-seq analysis supported the developmental direction of the spatial transcriptional signatures, direct multimodal measurements from matched specimens will be required to establish a more complete correspondence between morphological dynamics and molecular state. Future work should expand validation to additional microscopes, labeling strategies, organisms, and biological structures, incorporate uncertainty estimation and active-learning strategies to guide efficient human correction, and extend morphology-guided integration to matched longitudinal and molecular measurements across additional developmental systems.

Despite these limitations, FOCUS-3D establishes a data, model, and software foundation for general-purpose 3D cell segmentation in volumetric microscopy. By combining large-scale data curation, pretrained volumetric representation learning, end-to-end model building, and downstream cell-resolved analysis, including integration of cellular morphology with molecular state, it provides a practical starting point for converting complex 3D microscopy images into quantitative cellular representations. We hope that the dataset and open-source tools introduced in this work will encourage the community to share additional annotations and benchmarks, thereby supporting continued progress toward more accurate, generalizable, and user-adaptable 3D bioimage analysis.

## Acknowledgments

We would like to acknowledge the State Key Laboratory of Membrane Biology for support with confocal microscopy imaging. We also acknowledge support from the Imaging Core Facility, Technology Center for Protein Sciences, Tsinghua University for providing instrumentation and technical assistance. B.L. was supported by the Advanced Innovation Fellow Program of the Beijing Frontier Research Center for Biological Structure. We would like to thank the Zebrafish Information Network (ZFIN), FlyBase, WormBase, XenBase, and all the members of the Alliance of Genome Resources (Alliance) Consortium for their invaluable work and support of model organism research.

## Author contributions

Conceptualization, G.Y.; methodology, Q.Z., Z.M., B.L., J.N.A., Y.W., G.Y.; data, B.L., D.L., W.W., J.N., Y.W., J.N.A.; software, Q.Z.; evaluation, Q.Z., Z.M., Y.C.; writing, Q.Z., Z.M., B.L., Y.W., J.N.A. and G.Y.

## Code and data availability

- The code for FOCUS-3D is available on GitHub in Python at https://github.com/yu-lab-vt/FOCUS-3D.
- The website of FOCUS-3D is available at https://www.quiclab.org.cn/focus-3d.
- The gold-standard dataset of FOCUS-3D is being prepared for public release.

## Supplementary Notes

### Dataset collection

We collected and curated a large-scale 3D cell microscopy dataset from both public repositories and in-house sources. The complete dataset contains 47 sub-datasets, more than 17,000 image volumes and approximately 5.1 TB of data. The public datasets were collected from BBBC, Zenodo, BioImage Archive, Allen Cell Explorer, Cell Tracking Challenge, and IDR. These resources provide diverse 3D microscopy data, including synthetic 3D colon tissue with different SNR levels, *C. elegans* nuclei, mouse embryo nuclear and membrane data, 3D+time cell tracking benchmarks, and other large-scale published bioimage datasets. The in-house datasets include zebrafish, *Drosophila*, and mouse embryo imaging data. These datasets contain nuclear, membrane and cytoplasmic signals across different developmental stages and imaging conditions. In particular, the zebrafish datasets provide diverse embryonic patterns. This combination of public and self-collected data provides broad coverage of species, imaging modalities, labeling strategies, and cellular morphologies, supporting the development of a generalizable 3D cell instance segmentation model.

### Data organization

We organized the annotated data into two quality levels: silver-standard and gold-standard annotations. Silver-standard annotations were used for large-scale training and intermediate supervision, whereas gold-standard annotations were used for final fine-tuning and segmentation evaluation.

For public datasets, annotation quality was determined according to the reported annotation protocol and our manual inspection. We classified a public annotated dataset as silver-standard if it satisfied any of the following criteria: (i) the annotation was explicitly generated by a machine learning model, automatic segmentation algorithm, or semi-automatic computational method; (ii) the dataset was synthetic or simulated; (iii) visual inspection revealed clear annotation errors, such as missed cells, false positives, merged cells, fragmented instances, or inconsistent boundaries. Public annotations were not re-curated voxel-by-voxel; they were included as gold-standard annotations only when the original source reported manual annotation and our inspection did not reveal systematic errors.

For in-house datasets, we first generated silver-standard annotations using PrinCut-Auto^36^, an unsupervised 3D cell segmentation method. PrinCut-Auto is based on multi-scale principal curvature (MSPC), which enhances structural cues related to intercellular gaps and cell boundaries in dense 3D images. It further combines min-cut optimization with order-statistics testing to detect cellular structures. The PrinCut-Auto outputs were then manually reviewed and corrected by annotation experts. During expert curation, annotators corrected missing cells, false-positive detections, merged instances, over-segmented fragments, boundary errors, and slice-to-slice label inconsistencies. The complete correction process took 2,002 hours and included 321,734 editing operations. The resulting expert-corrected labels were used as the gold-standard annotations for in-house data.

### Benchmarking setup

In the benchmarking of peer methods, we evaluated performance before and after fine-tuning, as well as the slice-by-slice 2D segmentation performance of CellSAM. Inference time was measured on an NVIDIA A100 GPU with 80 GB of memory while constraining GPU memory usage to within 12 GB. For the fine-tuning comparison, all methods used the same training and testing sets. The zebrafish embryo data comprised a mixture of datasets 44 and 53, whereas the *Drosophila* and *C. elegans* embryo data were derived from datasets 2 and 19, respectively (see Table S1). For the comparison with slice-by-slice CellSAM, the zebrafish benchmark used dataset 44 only. All three-dimensional tuples are reported in (Z, Y, X) order.

#### FOCUS-3D

For FOCUS-3D inference, input volumes were processed using overlapping patches of size (32, 96, 96) with a stride of (24, 64, 64). The lower and upper percentile parameters for intensity normalization were set to 1 and 99, respectively. Query predictions were retained using a confidence threshold of 0.7 and a mask-probability threshold of 0.5. Connected components smaller than 64 pixels on individual *Z* slices were removed during post-processing. Inference was performed with a batch size of 8. The axial resolution ratio and cell-radius parameter were adjusted according to the imaging resolution and apparent cell size of each dataset.

#### Robustness evaluation

To evaluate robustness to common microscopy image perturbations, we independently tested image noise, lateral-resolution degradation, and intensity variation. For noise perturbations, images were first normalized to [0, 1] using the 0.1 and 99.9 intensity percentiles. Gaussian noise was added with standard deviations *σ* = 0.03, 0.1, 0.2, and 0.3. Poisson noise was simulated using maximum photon levels of 100, 10, 5, and 2, respectively. Mixed-noise conditions combined the corresponding Gaussian and Poisson settings (*σ, λ*_max_) = (0.03, 100), (0.1, 10), (0.2, 5), and (0.3, 2). Perturbed intensities were clipped to [0, 1] and mapped back to the original intensity range. To simulate reduced lateral resolution, images were downsampled in the X-Y plane by factors of 1.5, 2, 3, and 4 and subsequently upsampled to the original dimensions using linear interpolation. For intensity perturbations, linear intensity scaling factors of 0.25, 0.5, 1.5, and 2 were tested. Nonlinear intensity variations were generated by gamma transformations with *γ* = 0.5, 0.8, 1.5, and 2.0 after normalization to [0, 1].

#### StarDist

StarDist^26^ is a deep learning-based instance segmentation method that detects objects by predicting object probabilities together with a star-convex shape representation. In 2D, each object is represented by a star-convex polygon parameterized by radial distances from the object center; in 3D, this formulation is extended to star-convex polyhedra. This allows StarDist to directly produce instance-level segmentations for densely packed nuclei or cells in volumetric microscopy images. To achieve better performance within limited GPU memory, we tested various tiling parameters: (1, 2, 2) for *C. elegans* embryo, (1, 4, 4) for *Drosophila* embryo, and (4, 4, 4) for zebrafish embryo. During fine-tuning, the training configurations were set as follows: 50 epochs, 100 steps per epoch, and a batch size of 6 for zebrafish embryo and 24 for both *C. elegans* and *Drosophila* embryos. The training patch size was set to (32, 192, 192) for *C. elegans*, (48, 192, 192) for *Drosophila* embryo, and (32, 96, 96) for zebrafish embryo, with a learning rate of 3 *×* 10*^−^*^4^.

#### Cellpose

We used Cellpose-SAM, the SAM-based implementation of the Cellpose framework, for all Cellpose experiments in this study^13,44^; for simplicity, we refer to it as Cellpose throughout the text and figures. Cellpose represents cellular objects using a cell-probability map and spatial flow fields that guide pixels toward their corresponding object centers. Cellpose-SAM retains this flow-based segmentation formulation while incorporating a pretrained SAM transformer backbone to improve generalization. For 3D microscopy data, we applied the model in its native 3D inference mode, which estimates flows from orthogonal views of the volumetric stack and reconstructs 3D instance masks. For fine-tuning, we used 100 epochs, a learning rate of 1 *×* 10*^−^*^5^, a weight decay of 0.1, and a batch size of 24.

#### CellSAM

CellSAM^16^ is a foundation-model-based cell segmentation method adapted from the Segment Anything framework for microscopy cell images. It detects candidate cells by predicting bounding boxes and then generates instance masks using a SAM-based mask decoder. Since CellSAM does not provide training code for fine-tuning, we evaluated it only in the zero-shot setting. As CellSAM shows strong performance in 2D cell segmentation, we applied it slice-by-slice along the X-Y plane and evaluated its performance using 2D segmentation metrics. During inference, each 2D slice was normalized and processed in batches with batch size = 3. The bounding-box threshold was set to 0.4. The predicted masks were further post-processed by filling holes, removing small masks with a minimum size of 25 pixels, and subtracting object boundaries.

### Evaluation metrics

At a given IoU threshold *τ*, predicted–ground-truth pairs with IoU *≥ τ* were greedily matched in descending order of IoU, with each predicted and ground-truth instance allowed to be matched at most once. The numbers of true positives, false positives, and false negatives were then defined as the number of matched pairs, unmatched predictions, and unmatched ground-truth instances, respectively. Unless otherwise specified, F1 scores were calculated at IoU = 0.5. Precision, recall, and F1 score were computed as

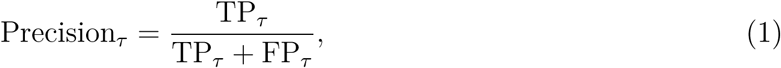

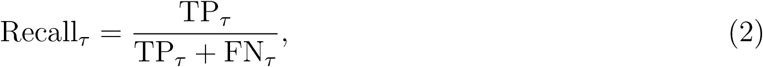

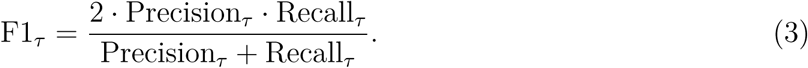

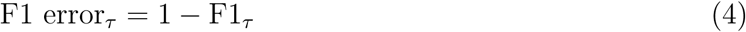

We reported AP@0.5 and AP@0.75 using the object-level definition

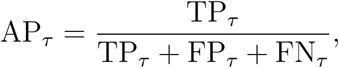

following the metric form used in Cellpose.

### Network architecture

Our proposed network follows a Mask2Former-style query-based mask classification paradigm^29^ and is adapted to volumetric instance segmentation. It consists of a 3D MAE backbone^37^, an anisotropy-aware ViT-Adapter^41^, a 3D multi-scale deformable attention pixel decoder, and a masked transformer decoder^29^ (Figure 2).

#### 3D MAE backbone

The backbone extracts volumetric semantic features from input patches and provides a generalizable representation learned from large-scale unlabeled 3D microscopy data. The input volume size is set to 32×96×96, with a patch size of 8×12×12. The MAE encoder uses an embedding dimension of 1008, 24 transformer layers, 16 attention heads, and an MLP ratio of 4.0. A decoder with embedding dimension 768 and 8 transformer layers is used during MAE pretraining. This design replaces a conventional supervised segmentation backbone with a self-supervised 3D representation backbone, improving robustness across different cell types, imaging modalities, and annotation styles.

#### ViT-Adapter

The ViT-Adapter converts the single-scale MAE transformer representation into hierarchical feature maps suitable for dense prediction. Four interaction stages are used, corresponding to MAE encoder layers [0, 5], [6, 11], [12, 17], and [18, 23]. To better handle anisotropic microscopy volumes, the early feature pyramid stages avoid downsampling along the Z axis, preserving axial information when the axial resolution is substantially lower than the lateral resolution.

#### 3D pixel decoder

The pixel decoder aggregates multi-scale volumetric features and produces high-resolution mask features for instance mask prediction. The convolution dimension and mask feature dimension are both set to 288, and the transformer encoder contains 6 layers. Compared with a 2D pixel decoder, all convolution, attention, and feature-fusion operations are extended to 3D, allowing spatial context to be integrated jointly across three dimensions.

#### Masked transformer decoder

The transformer decoder predicts instance-level object queries and converts them into 3D masks through a mask head. It uses 300 object queries, hidden dimension 288, 8 attention heads, feed-forward dimension 2048, and 9 decoder layers. The training objective combines classification, mask, and Dice losses with weights 2.0, 5.0, and 5.0, respectively, and the no-object class weight is set to 0.1.

Two targeted training strategies are further introduced in the decoder. First, denoising auxiliary training is used to stabilize query learning, where 100 denoising queries are generated with segmentation-style perturbations, with a denoising loss weight of 1.0. Second, object query initialization is performed from encoded volumetric features rather than relying solely on randomly learned queries. Candidate object proposals are ranked from the encoded feature map, and the top-ranked proposals are used to initialize query content and query position. This encourages the decoder queries to start from object-aware 3D locations, improving convergence and reducing duplicate or background-dominated queries.

### Inference

For full-volume segmentation, inference was performed using overlapping 3D blocks. Each non-background block (size 32 *×* 96 *×* 96) was processed independently by the trained model, and the query-level predictions were converted into a local instance map. The local results were then stitched into a global instance map, followed by a final filtering step to remove small or low-confidence fragments.

#### Within-block post-processing

For each block, the model predicted a set of query masks and corresponding confidence scores. We first computed a mask-quality score for each query as the average mask probability over voxels above the mask threshold. The final query confidence was defined as the product of the object score and the mask-quality score. Queries were retained only if they passed both the confidence and mask-quality thresholds.

Due to the frequent high overlap of two cells’ X-Y projections in adjacent *z*-slices under anisotropic imaging, where query predictions sometimes produce overlapping masks at the interface *z*-planes, we resolved local conflicts between these overlapping query masks before assigning voxels to instances. On each *z*-slice, regions covered by multiple queries were detected. If two masks strongly overlapped, measured by IoU and IoM, they were treated as representing the same local structure. The merged region was assigned to the query that showed better consistency with its neighboring *z*-slices. Other overlapping queries were suppressed only within the merged region, while their non-overlapping regions were preserved.

After conflict resolution, each foreground voxel was assigned to the query with the highest confidence-weighted mask probability:

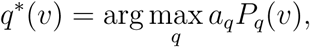

where *P_q_*(*v*) is the mask probability of query *q* at voxel *v*, and *a_q_* is the final query confidence. Voxels whose selected mask probability was below the mask threshold were set to background. This produced a local instance map and a corresponding local confidence map for each block.

#### Block stitching

The local instance maps were stitched into a global instance map during inference. For each incoming block, local instance IDs were compared with existing global IDs along the neighboring block faces. If a local instance overlapped sufficiently with an existing global instance on the contact faces, the local ID was remapped to the matched global ID. Otherwise, a new global instance ID was assigned. This face-based matching allowed objects crossing block boundaries to be connected while avoiding unnecessary merging of unrelated instances.

In overlapping regions, voxel confidence values from multiple blocks were accumulated and averaged:

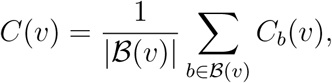

where *B*(*v*) denotes the set of blocks contributing to voxel *v*, and *C_b_*(*v*) is the confidence assigned by block *b*. This yielded a stitched global instance map and a voxel-wise confidence map.

#### Final post-processing

After stitching, final post-processing was applied to remove fragmented predictions. First, on each *z*-slice, components smaller than the area threshold were removed from both the instance map and the confidence map. Second, 3D instances that did not satisfy the size or intensity criteria were removed. The final output consisted of a 3D instance map and a corresponding confidence map.

### Training details

Before supervised training, all 3D volumes were cropped into fixed-size training blocks of 32 *×* 96 *×* 96 voxels. To mitigate batch effects and improve generalization across different imaging resolutions and cell sizes, we first normalized the annotated cells in the ground truth to a unified diameter of 30 voxels. Subsequently, during block generation, we applied random resizing to these normalized volumes to deliberately simulate variations in apparent cell scale. Blocks that contained no annotated cells were excluded from the training set.

During supervised training, we applied online data augmentation to improve generalization across diverse 3D microscopy datasets. Geometric augmentations included random flipping along the Z, Y, and X axes and random 90-degree rotations in the X-Y plane. Photometric augmentations included random brightness and contrast perturbations. To mimic common microscopy degradations, we further applied random image-only degradation with a fixed probability, including Poisson noise, Gaussian blur, downsampling followed by upsampling, and anisotropic blur. After augmentation, image intensities were clipped to the valid normalized range.

The model was trained in three stages. First, we performed self-supervised 3D MAE pretraining using unlabeled volumetric image blocks. The MAE was trained for 400 epochs with AdamW optimization and a weight decay of 0.05. Second, the pretrained backbone was used to initialize the FOCUS-3D segmentation model, which was trained on silver-standard annotations for 80,000 iterations with a learning rate of 1*×*10*^−^*^4^. Third, the model was further fine-tuned on gold-standard annotations for 40,000 iterations with a reduced learning rate of 5 *×* 10*^−^*^5^.

### Interactive curation platform

#### User interface and interactive curation

The annotation interface was designed to support user-friendly 3D instance curation and human-in-the-loop model improvement (Figure 4, Figure S6). Basic curation operations include adding voxels to an existing label, subtracting voxels from a label, creating a new label, deleting selected labels, and deleting labels within a user-defined ROI. The interface also supports block-wise annotation for large volumes, one-click segmentation using the pretrained model, fine-tuning with newly curated labels, quantitative measurement, 3D instance reconstruction, and dual-channel visualization. These functions allow users to iteratively correct segmentation results, generate high-quality annotations, and update the model within the same workflow.

#### One-click segmentation

For interactive correction, we implemented a one-click instance inference mode. Given a user-selected voxel, a local 3D patch centered at the clicked position was extracted and passed through the trained model (Figure 4B). The model produced a set of query-level object scores and mask probability maps. To identify the clicked object, each query was evaluated according to its mask response in a small neighborhood around the clicked voxel, rather than at a single voxel, making the selection more robust to boundary clicks and weak local signals. Only the connected component containing the clicked voxel was retained. The resulting local mask was then mapped back to the original image coordinates as the one-click instance mask.

### Experiments

#### Animal model and subject details

Wild-type (WT) zebrafish of the Tuebingen (Tu) strain were used in this study. Adult fish were maintained in a recirculating water system at 28.5 °C. After fertilization, embryos were incubated at 28.5 *^◦^*C in Holtfreter’s solution (0.059 M NaCl, 0.00067 M KCl, 0.00076 M CaCl_2_, and 0.0024 M NaHCO_3_). All experimental procedures involving zebrafish embryos were approved by the Animal Care and Use Committee of Tsinghua University and performed in accordance with their guidelines.

#### Live imaging preparatio

To label cellular membranes and nuclei for live imaging, one-cell stage WT embryos were injected with 100 pg of eGFP-CAAX mRNA (membrane marker) and 100 pg of NLS-mCherry mRNA (nuclear marker). Injected embryos were allowed to develop to 10 hours post-fertilization (hpf). Prior to imaging, the chorions were removed by treatment with Pronase (protease from Streptomyces griseus, Sigma, cat. no. P5147). Dechorionated embryos were then embedded in 1 % low-melting point agarose.

#### Plasmid construction and RNA synthesis

For plasmid construction, the eGFP and CAAX sequences, as well as the NLS and mCherry sequences, were assembled into the pCS2+ vector by homologous recombination-based cloning (Lablead, D0204P) to generate the eGFP-CAAX and NLS-mCherry constructs, respectively. All recombinant plasmids were verified by DNA sequencing prior to in vitro transcription.

For mRNA synthesis, plasmids were linearized with NotI-HF (New England Biolabs, R3189V) at 37 *^◦^*C, followed by heat inactivation of the restriction enzyme. The linearized DNA was purified using the Universal DNA Purification Kit (TIANGEN, DP214). RNA was synthesized from the purified linearized templates using the mMESSAGE mMACHINE^™^ SP6 Transcription Kit (Thermo Fisher Scientific, AM1340) according to the manufacturer’s instructions. The resulting mRNA was purified using VAHTS RNA Clean Beads (Vazyme, N412-01) according to the manufacturer’s protocol and eluted in nuclease-free water. RNA concentration was measured using a NanoDrop spectrophotometer. Purified mRNA was aliquoted and stored at *−*80 *^◦^*C until use.

#### Image acquisition and processing

Time-lapse z-stack imaging was performed on a spinning-disk confocal microscope using a 40× objective. Z-stacks were acquired with a step size of 1 µm, and the time interval between frames was 1.5 min. Images were processed using Imaris software (version 8.1.4, Bitplane AG).

### Morphometric, positional and dynamic feature definitions

All single-cell morphometric and dynamic features were computed in physical coordinates using the dataset-specific voxel spacing. For voxel spacing (*s_z_, s_y_, s_x_*), a voxel coordinate (*z, y, x*) was represented as

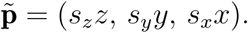

For a cell mask containing *N* foreground voxels, cell volume was calculated as

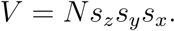

Surface area, denoted as *A*, was estimated from the triangular surface mesh reconstructed from the 3D binary mask using marching cubes with the corresponding dataset-specific voxel spacing.

Cell sphericity was calculated as

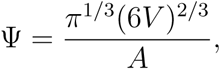

where Ψ = 1 corresponds to a perfect sphere and lower values indicate increasing deviation from spherical geometry.

To characterize 3D cell shape, effective ellipsoidal axis lengths were estimated from the eigenvalues of the 3D inertia tensor. After sorting the eigenvalues as *λ*_1_ *≥ λ*_2_ *≥ λ*_3_, the major, intermediate, and minor axis lengths were calculated as

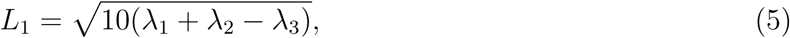

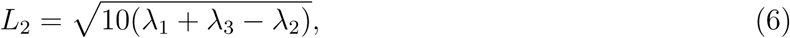

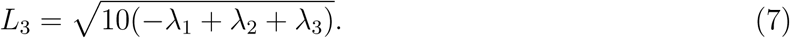

Three complementary axis-ratio descriptors were then defined as

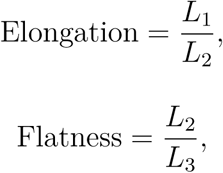

and

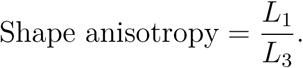

Higher elongation indicates preferential extension along the major axis, whereas higher flatness reflects a stronger difference between the intermediate and minor axes. Shape anisotropy summarizes the overall disparity between the longest and shortest cell dimensions.

Cell extent was defined as the fraction of the 3D bounding-box volume occupied by the segmented cell,

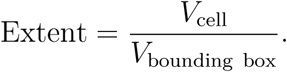

The maximum-section axis ratio was computed from the 2D slice containing the largest cell cross-sectional area. Foreground pixels in this slice were subjected to 2D principal-axis analysis, and the ratio between the long and short axes was reported as

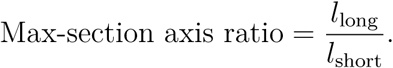

Cell movement was quantified from the displacement of the cell centroid between consecutive analyzed time points. For a tracked cell with centroid (*z_t_, y_t_, x_t_*) at time *t*, the anisotropy-corrected centroid was

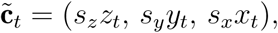

and movement over one time interval was calculated as

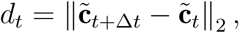

where Δ*t* denotes the interval between adjacent analyzed frames.

For comparisons between the lineage-defined notochord population and surrounding cells, the relative difference of a morphometric feature *X* at time *t* was calculated as

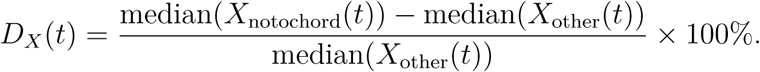

Positive values indicate higher feature values in notochord cells, whereas negative values indicate lower values relative to surrounding cells.

#### Definition of the 6-somite AP coordinate

The longitudinal coordinate of the 6-somite notochord was defined directly from physical centroids of the corresponding instances. Principal-component analysis was applied to the 579 notochord cell centroids, and the first principal component was used as the longitudinal axis. Axis orientation was determined from the annotated Notochord Anterior and Notochord Posterior Tip–posterior populations such that increasing values pointed toward the posterior end. Projected cell coordinates were then linearly normalized to the interval [0, 1], with 0 corresponding to the anterior end and 1 to the posterior end.

#### Feature normalization and morphology similarity

Seven morphometric features were used for cross-dataset morphology matching: cell volume, surface area, sphericity, elongation, flatness, shape anisotropy, and extent. Before normalization, volume and surface area were transformed using log(1+*x*), whereas elongation, flatness, and shape anisotropy were log-transformed. Morphological features were then standardized independently within the live-imaging and weMERFISH datasets using robust scaling,

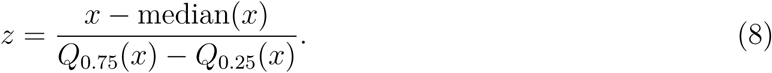

If the interquartile range was zero or undefined, the standard deviation was used instead. The seven standardized morphology features were assigned equal weights of 1/7.

For temporal state *t* and spatial window *w*, feature-specific dissimilarity was calculated using the one-dimensional Wasserstein distance,

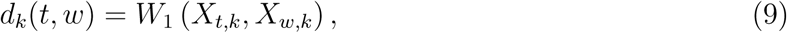

where *X_t,k_* and *X_w,k_* are the distributions of standardized morphology feature *k* in the temporal and spatial populations, respectively. Aggregate morphology similarity was defined as

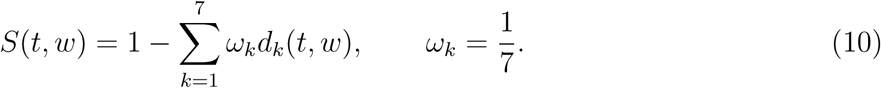

#### Morphology-based temporal-to-spatial mapping

Candidate spatial regions were generated by sliding a window spanning 10% of the normalized AP length along the 6-somite notochord. Window starting positions were evaluated at increments of 0.005. The window giving the maximum morphology similarity was selected independently for each temporal state. Intermediate frames were retained for visualization of the morphology-matching trajectory, whereas downstream transcriptional analyses compared only the two endpoint states, frame 0 and frame 80. The spatial support interval of each morphology match was defined from the continuous similarity peak surrounding the best-matching AP position. For each temporal state, the support threshold was calculated as

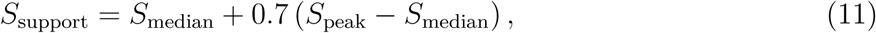

where *S*_peak_ is the maximum morphology-similarity score and *S*_median_ is the median similarity across all candidate AP windows. Starting from the global maximum, the contiguous AP interval remaining above this threshold was identified, with threshold crossings linearly interpolated between adjacent windows. This interval was used as the AP support range shown in Figure 5H.

With this configuration, the optimal frame 0 spatial analogue occupied AP 0.900–1.000 and contained 166 cells, with a morphology-similarity score of 0.346. The optimal frame 80 spatial analogue occupied AP 0.505–0.605 and contained 57 cells, with a similarity score of 0.616. The two regions contained no overlapping cells. The frame 0-like population consisted of 59 Notochord Posterior Tip–anterior cells and 107 Notochord Posterior Tip–posterior cells, whereas all 57 frame 80-like cells belonged to the Notochord Anterior population.

#### Spatial differential-expression analysis

The expression profiles of the morphology-defined frame 0-like and frame 80-like populations were compared using the 495 genes contained in the weMERFISH AnnData object. For each gene, a two-sided Mann–Whitney *U* test was performed between the 166 frame 0-like cells and 57 frame 80-like cells. Effect size was quantified using rank-biserial correlation,

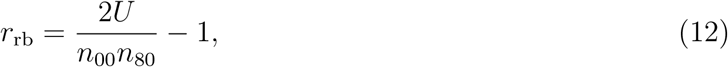

where positive values denote higher expression in the frame 0-like population and negative values denote higher expression in the frame 80-like population. Gene-wise *P* values were corrected using the Benjamini–Hochberg procedure. Genes with FDR *<* 0.05 were retained as spatially differential, without an additional effect-size threshold. This yielded 293 genes, including 109 genes higher in the frame 0-like region and 184 genes higher in the frame 80-like region.

#### Independent single-cell RNA-seq analysis

Independent developmental-stage validation was performed using cells annotated as notochord segment in the single-cell RNA-seq dataset. Cells from the bud stage (ZFB; *n* = 148) were compared with a combined somite-stage group consisting of 27 3-somite cells (ZF3S) and 70 6-somite cells (ZF6S), for a total of 97 somite-stage cells.

The raw expression matrix was used as the expression source. Counts were normalized to 10^4^ total counts per cell and transformed using log(1 + *x*). Differential expression between the combined ZF3S+ZF6S group and ZFB was calculated using Scanpy’s Wilcoxon rank-sum test. Positive log_2_ fold changes indicate higher expression in ZF3S+ZF6S, whereas negative values indicate higher expression in ZFB. Genes were classified as temporally differential when

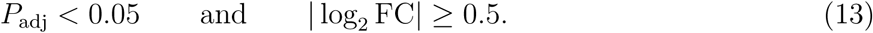

For genome-wide characterization of developmental transcriptional differences, all 17,239 genes in the single-cell RNA-seq matrix were tested. This analysis identified 157 differentially expressed genes, including 129 genes with higher expression at the bud stage and 28 genes with higher expression in the combined ZF3S+ZF6S group. These genome-wide differential-expression results were used for Gene Ontology analysis.

For direct cross-dataset concordance analysis, the single-cell RNA-seq data were separately restricted to genes measured by weMERFISH. The weMERFISH panel file contained 495 unique gene names after duplicate removal, of which 456 could be matched to the single-cell RNA-seq matrix. No additional minimum-expression filtering was applied. Among these 456 genes, 45 met the differential-expression criteria, including 35 genes higher at the bud stage and 10 genes higher at somite stages.

#### Gene Ontology enrichment analysis

Gene Ontology (GO) enrichment analysis was performed in R using topGO with *Danio rerio* annotations from org.Dr.eg.db. Gene symbols were first mapped to Ensembl gene identifiers using a local Ensembl release 100/Zv11 mapping table. Genes that could not be mapped in this table were additionally matched using the SYMBOL and ALIAS annotations in org.Dr.eg.db. Only Ensembl identifiers represented in org.Dr.eg.db were retained for enrichment analysis.

For the weMERFISH analysis, genes expressed more highly in the frame 0-like and frame 80-like regions were analyzed separately. The 495 genes measured in the weMERFISH dataset were used as the source background universe, and the effective enrichment background consisted of genes from this panel that could be mapped to Ensembl identifiers and annotated in org.Dr.eg.db. GO Biological Process enrichment was performed using the weight01 algorithm with Fisher’s exact test and a minimum node size of 5 annotated genes.

For the independent single-cell RNA-seq analysis, the 129 genes expressed more highly at the bud stage and the 28 genes expressed more highly in the combined ZF3S+ZF6S group were analyzed separately. The 17,239 genes tested in the genome-wide differential-expression analysis were used as the source background universe, with the effective enrichment background restricted to successfully mapped and annotated Ensembl genes. GO Biological Process enrichment was performed using the weight01 algorithm with Fisher’s exact test and a minimum node size of 10 annotated genes.

For both analyses, the enrichment factor was calculated as the observed number of selected genes annotated to a GO term divided by the expected number based on the corresponding effective background. Raw weight01 Fisher *P* values were used to rank GO terms, and terms with *P <* 0.05 were considered for visualization; up to the 15 most significant terms were displayed. Benjamini– Hochberg-adjusted *P* values were additionally calculated as a conservative sensitivity statistic. In the bubble plots, point size represents the number of selected genes annotated to each GO term and color represents *−* log_10_(*P* ) based on the raw weight01 Fisher test.

#### Cross-dataset concordance analysis

The 109 frame 0-associated and 184 frame 80-associated weMERFISH genes were matched to the 456-gene single-cell testing universe. This retained 99 frame 0-associated genes and 170 frame 80-associated genes. Directional overlap was evaluated using one-sided Fisher’s exact tests with the 456 tested genes as the background universe.

Four comparisons were evaluated: frame 0-associated genes versus bud-upregulated genes, frame 80-associated genes versus somite-stage-upregulated genes, and the two opposite-direction combinations as controls. Benjamini–Hochberg correction was applied jointly across the four Fisher tests. Frame 0-associated genes were significantly enriched among bud-stage-upregulated genes (19 overlaps; odds ratio = 5.06; *P* = 1.1 *×* 10*^−^*^5^; FDR = 4.2 *×* 10*^−^*^5^), and frame 80-associated genes were significantly enriched among somite-stage-upregulated genes (8 overlaps; odds ratio = 7.01; *P* = 6.97 *×* 10*^−^*^3^; FDR = 1.39 *×* 10*^−^*^2^). Neither cross-direction comparison was significant (frame 80 versus bud: 8 overlaps, odds ratio = 0.47, FDR = 0.981; frame 0 versus somite stage: 1 overlap, odds ratio = 0.39, FDR = 0.981).

## Supplementary Figures

**Figure S1:**
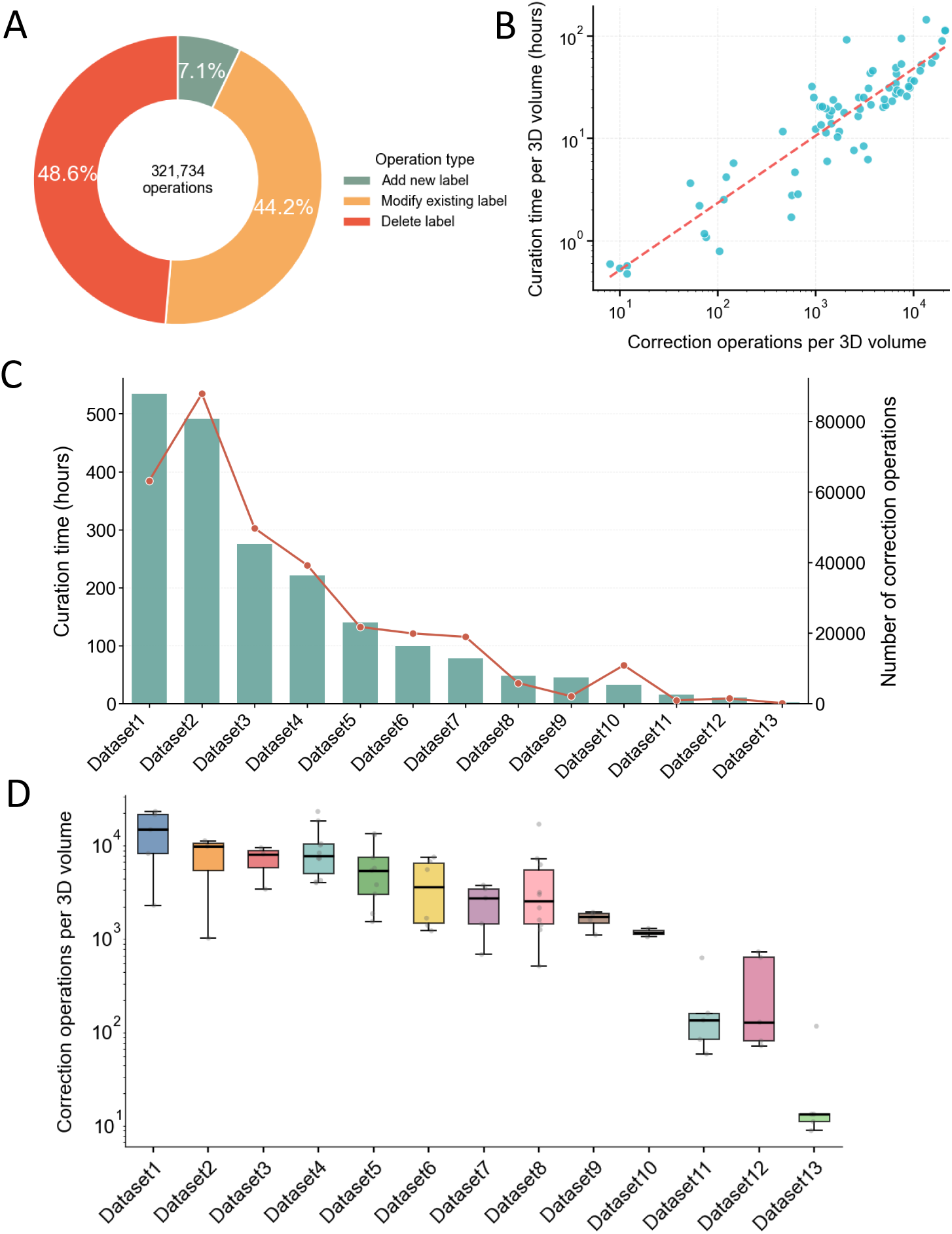
Quantification of manual curation workload across 3D microscopy datasets. (A) Distribution of correction operation types recorded during manual curation, showing the relative contribution of different editing actions. (B) Relationship between the number of correction operations per 3D volume and the corresponding active curation time, indicating that volumes requiring more editing generally involved longer manual curation. (C) Dataset-level summary of total active curation time and total number of correction operations across 13 datasets, highlighting the substantial manual effort required to generate high-quality 3D annotations. (D) Boxplot of correction operations per 3D volume across the 13 datasets, showing the variability of volume-level curation workload within and between datasets.

**Figure S2:**
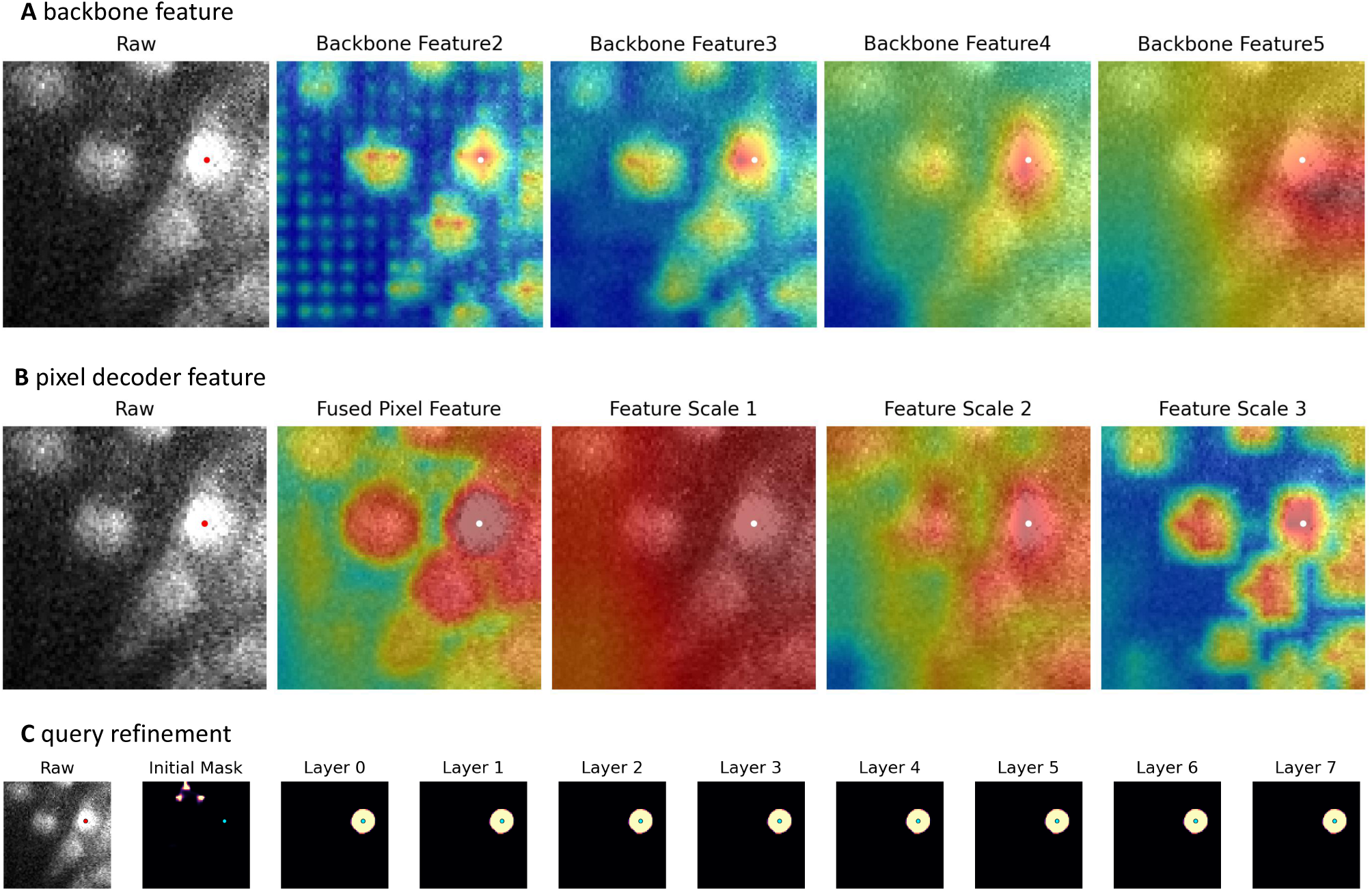
Visualization of intermediate representations and query refinement in the FOCUS-3D architecture. (A) Features extracted by the MAE backbone. A reference point was selected in the raw image, and the spatial correlations between the feature representation at this point and feature representations at other locations were visualized across different feature scales. (B) Features extracted by the pixel decoder. Using the same strategy, spatial correlations between the feature representation at a selected reference point and those at other locations were visualized across different decoder feature scales. (C) Query refinement across the transformer decoder. For a query associated with a selected location in the raw image, the predicted mask was visualized across successive decoder layers, illustrating the progressive refinement of the instance prediction.

**Figure S3:**
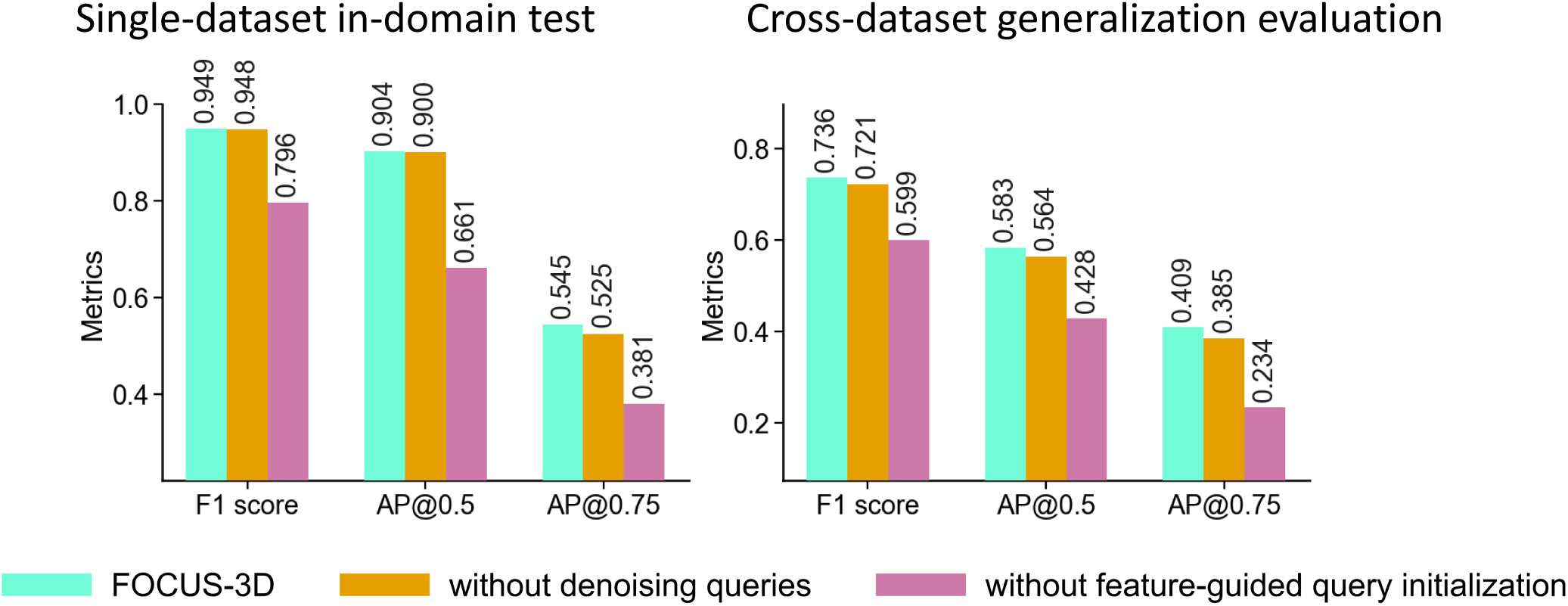
Ablation studies of the FOCUS-3D architectural innovations. We evaluated FOCUS-3D and its variants without denoising queries or without feature-guided query initialization under two settings: a single-dataset in-domain test, in which models were trained and evaluated on zebrafish embryo cell data from the same dataset, and a cross-dataset generalization evaluation, in which models were trained on a mixed training set consisting of zebrafish, *Drosophila*, and *C. elegans* embryo datasets and evaluated on zebrafish embryo data acquired under a different imaging condition.

**Figure S4:**
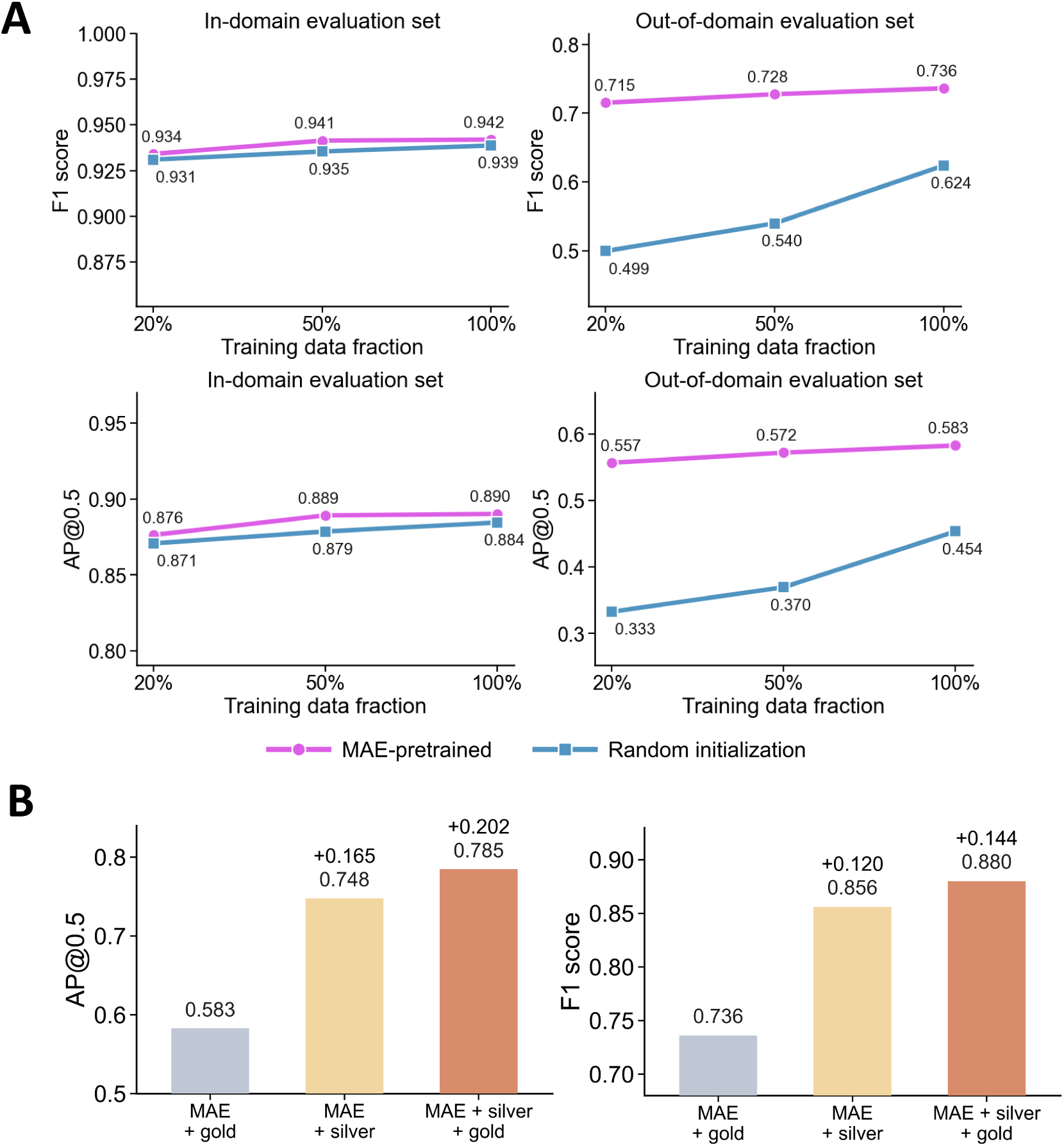
Ablation of the multistage training strategy. (A) Ablation of MAE initialization. Models were trained using 20%, 50%, or 100% of the mixed training set consisting of zebrafish, *Drosophila*, and *C. elegans* embryo datasets. We compared MAE-pretrained initialization with random initialization of the MAE parameters and evaluated both F1 score and AP@0.5 on two test settings: an in-domain zebrafish embryo test set from a dataset represented in training and a cross-dataset zebrafish embryo test set acquired under an unseen imaging condition. (B) Ablation of the silver-standard and gold-standard training stages. Starting from the same MAE-pretrained initialization, we compared three training configurations: gold-standard training alone (MAE + gold), silver-standard training alone (MAE + silver), and silver-standard training followed by gold-standard refinement (MAE + silver + gold). Segmentation performance was evaluated using AP@0.5 and F1 score on the same cross-dataset zebrafish embryo test set acquired under an unseen imaging condition.

**Figure S5:**
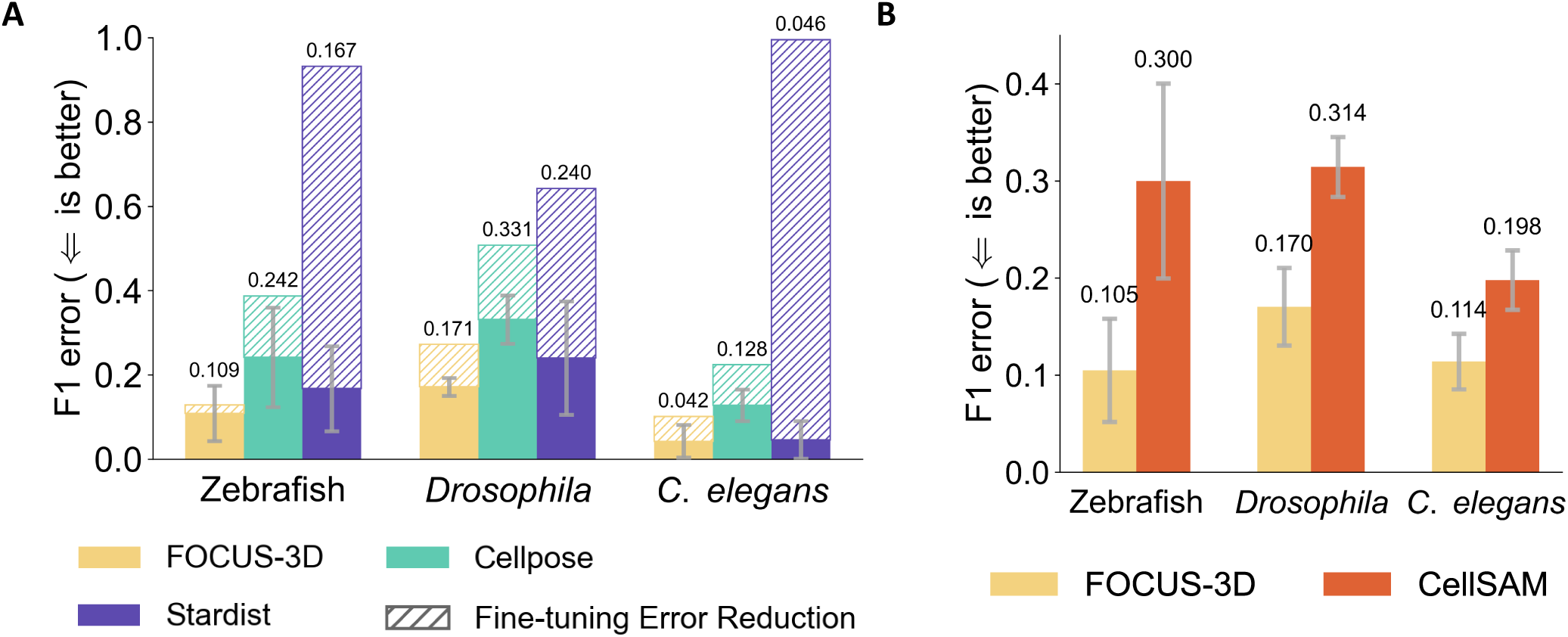
Segmentation benchmark of FOCUS-3D against competing methods. (A) Comparison of F1 error (1 − F1 score) before and after fine-tuning between FOCUS-3D, Cellpose, and StarDist across different datasets. FOCUS-3D achieves the best performance both before and after fine-tuning. (B) Comparison of F1 error between FOCUS-3D and the slice-by-slice 2D segmentation results of CellSAM.

**Figure S6:**
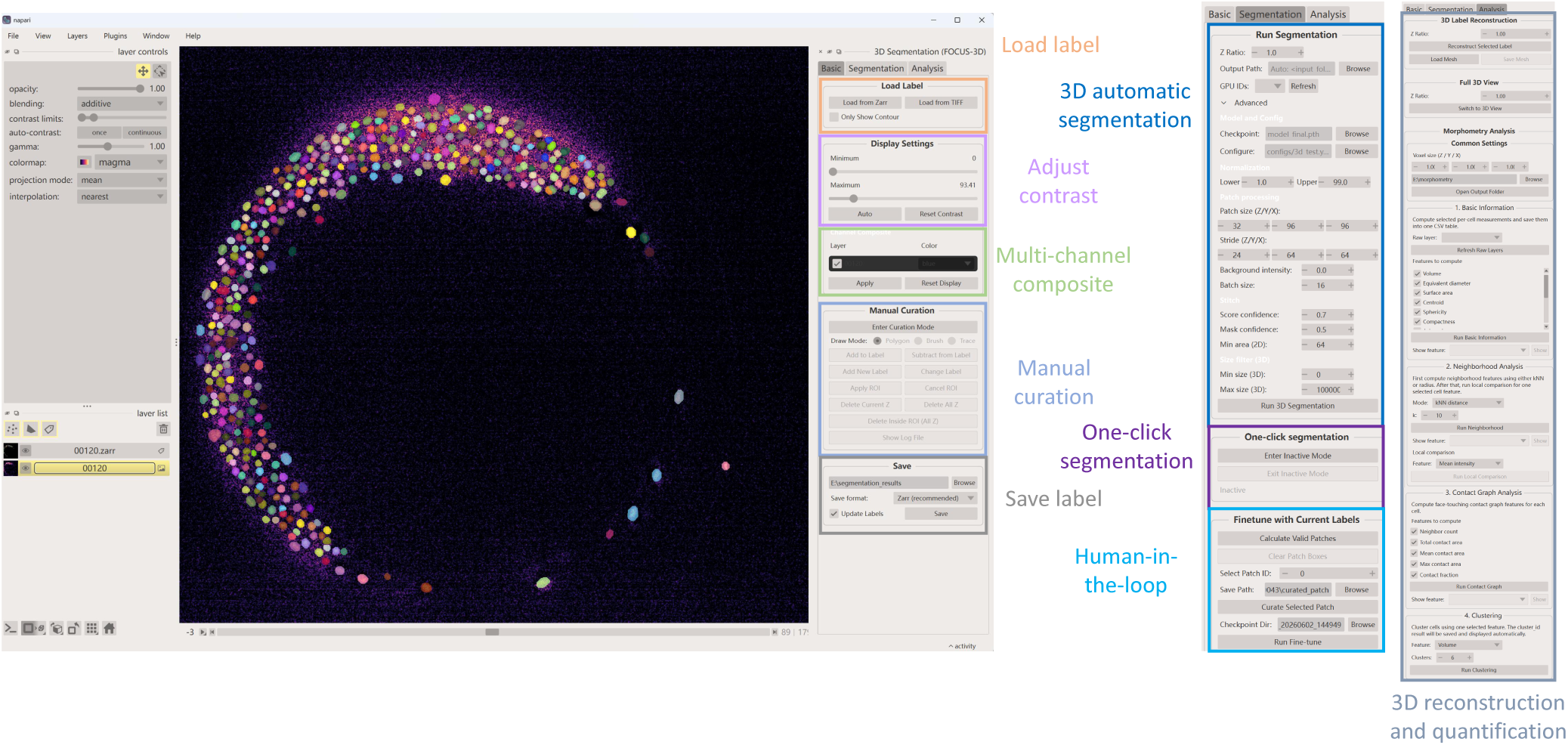
The napari plugin interface of FOCUS-3D. This interface integrates a complete pipeline of cell segmentation and downstream analysis.

**Figure S7:**
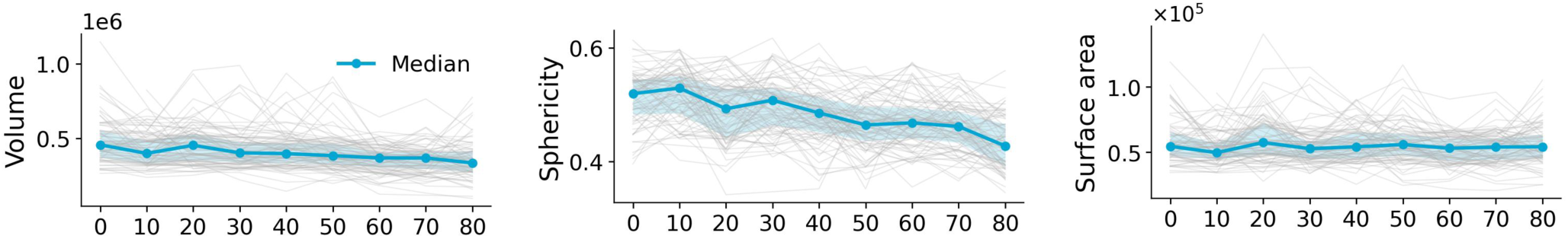
Temporal changes in single-cell anisotropy-corrected volume, sphericity and surface area within the selected notochord subregion. Gray lines represent individual tracked cells, and the blue line represents the median across cells.

**Figure S8:**
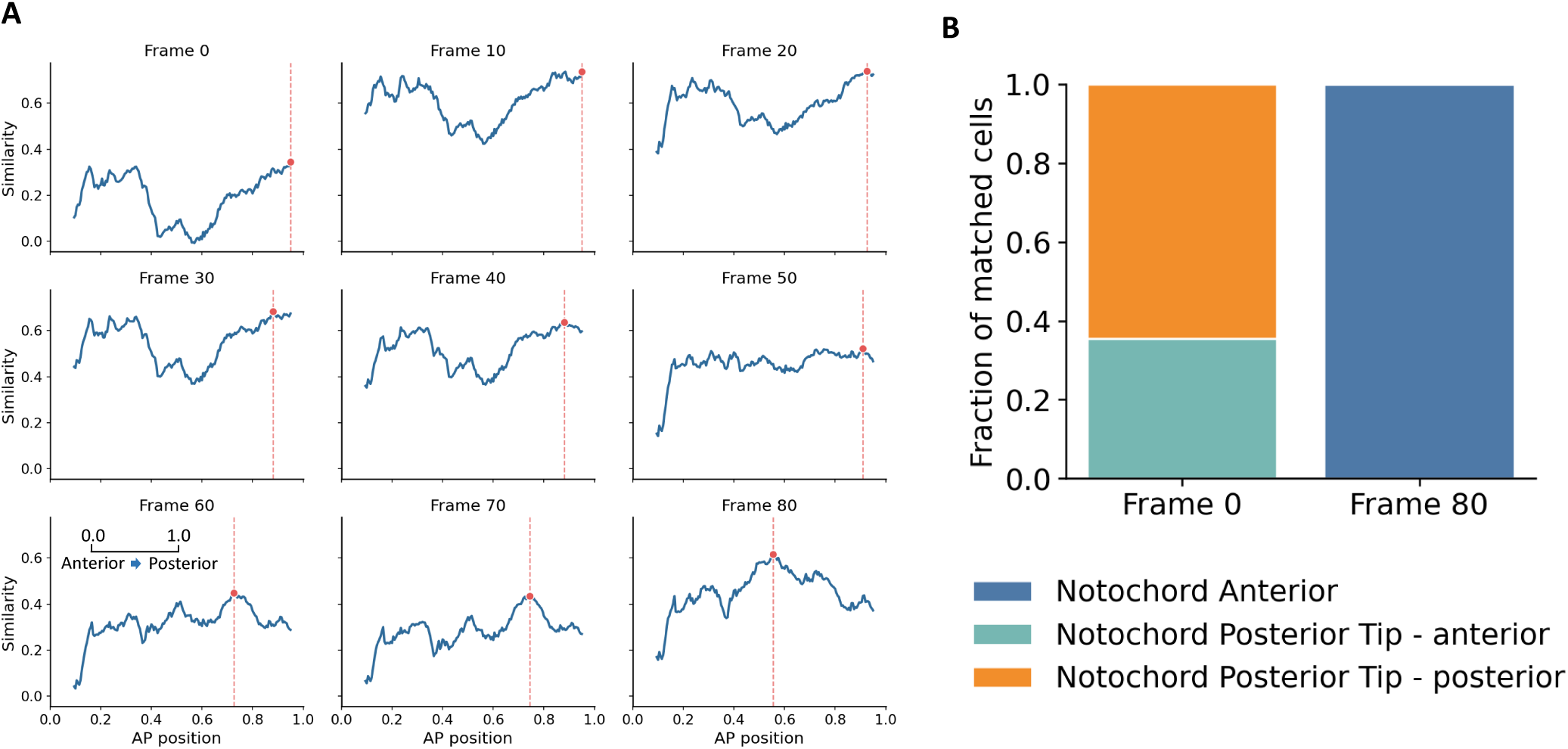
Morphology-guided temporal-to-spatial mapping of notochord cells. (A) Morphology-similarity landscapes between each temporal notochord state from frame 0 to frame 80 and 10% sliding windows along the anterior–posterior (AP) axis of the 6-somite weMERFISH notochord. The red point and dashed line indicate the spatial window with the highest morphology similarity for each time point. (B) Composition of the weMERFISH notochord cell populations within the morphology-matched regions for frame 0 and frame 80. Bars show the fraction of cells assigned to the Notochord Anterior, Notochord Posterior Tip–anterior, and Notochord Posterior Tip–posterior populations.

**Figure S9:**
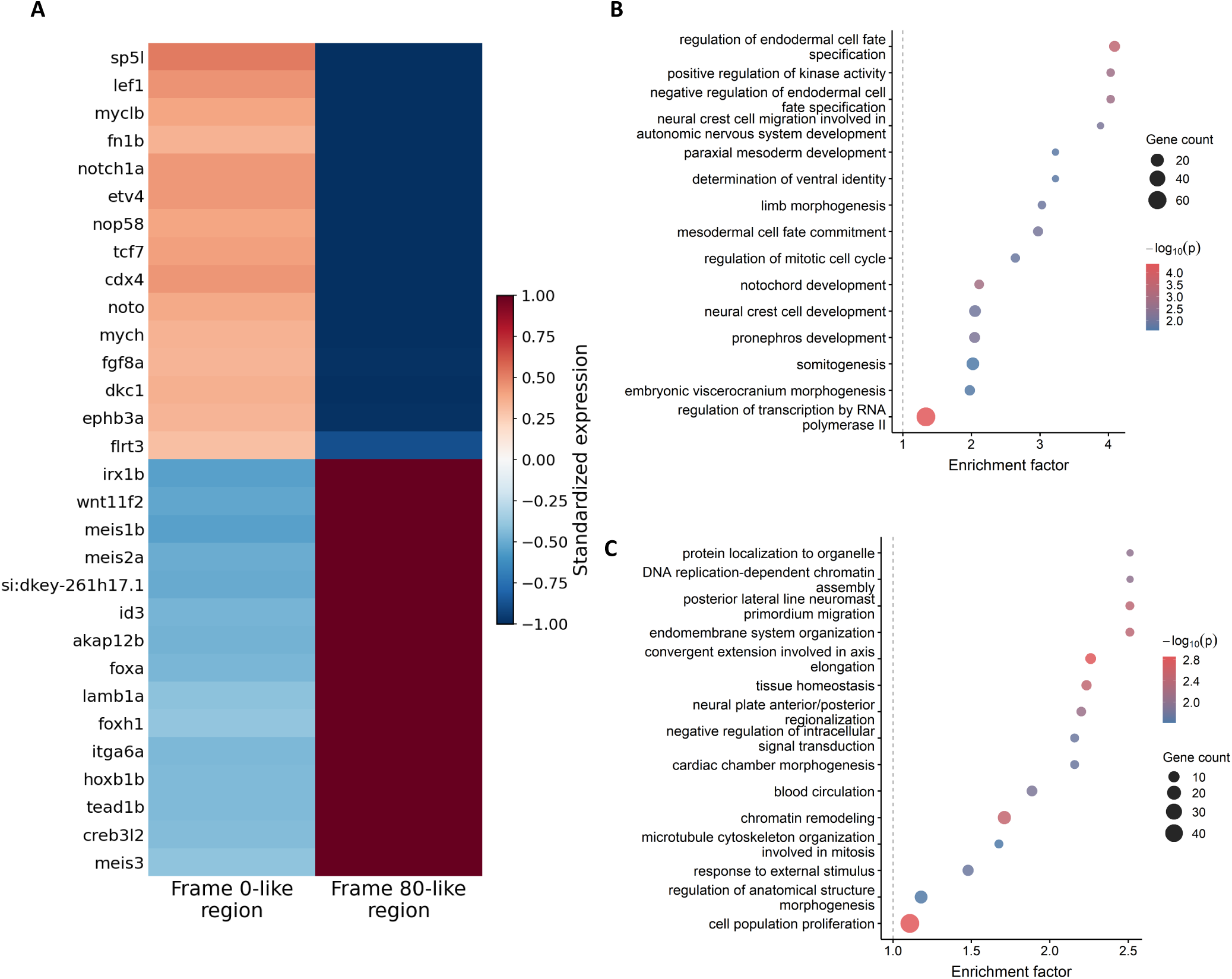
Transcriptional differences between morphology-matched frame 0-like and frame 80-like notochord regions. (A) Heatmap of representative differentially expressed genes between the frame 0-like and frame 80-like regions in the weMERFISH dataset. Genes were selected from the most significant genes enriched in either region, and expression was standardized across cells for visualization. (B,C) Gene Ontology biological-process enrichment analyses of genes expressed more highly in the frame 0-like and frame 80-like regions, respectively. Bubble size indicates gene count, color represents *−* log_10_(*P* ), and the x-axis shows the enrichment factor.

**Figure S10:**
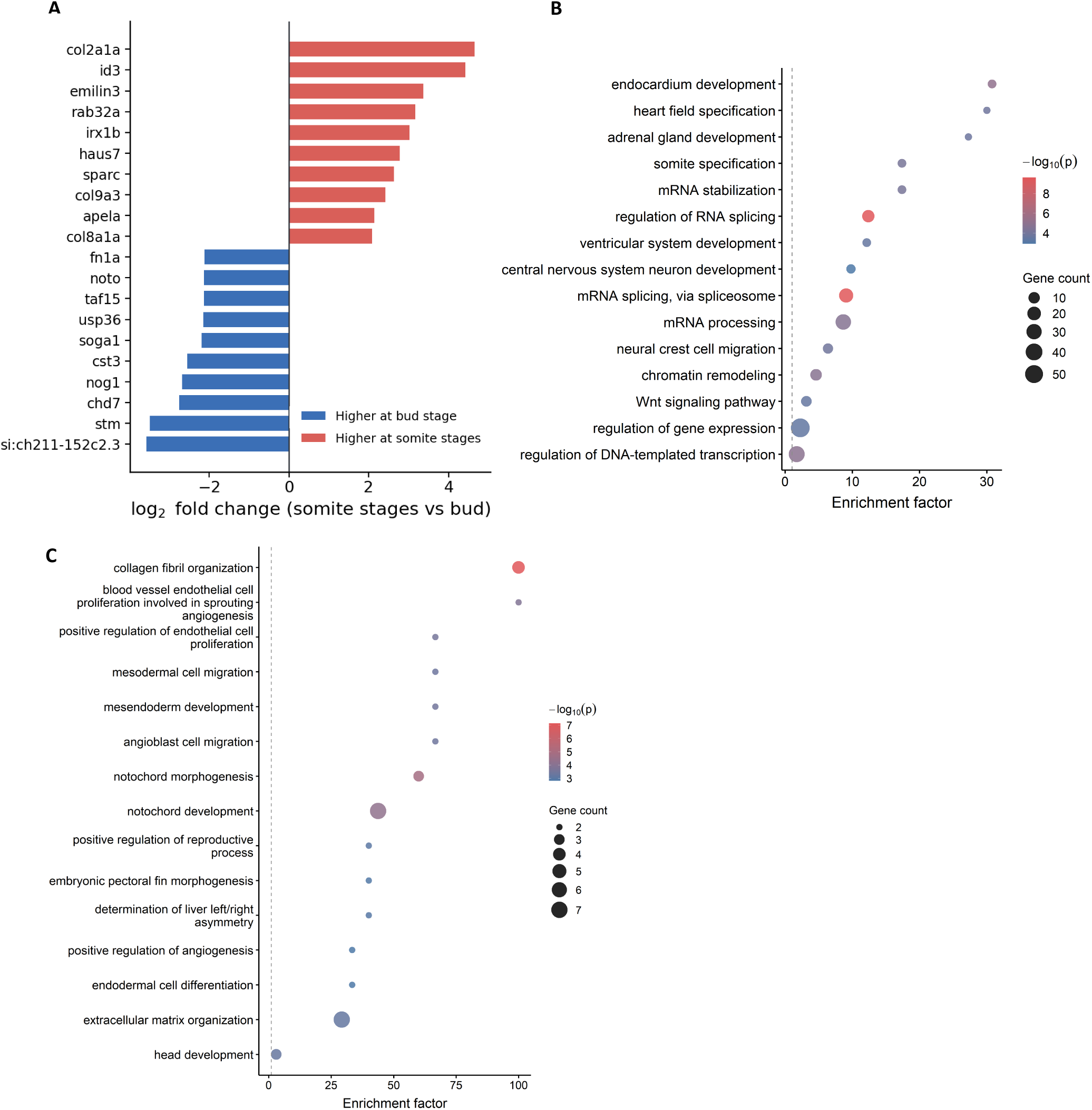
Developmental transcriptional differences in notochord cells identified by single-cell RNA sequencing. (A) Representative differentially expressed genes identified by full-transcriptome comparison of bud-stage notochord cells with the combined 3-somite/6-somite population. Negative log_2_ fold changes indicate higher expression at the bud stage and positive values indicate higher expression at somite stages. (B) Gene Ontology biological-process enrichment analysis of genes expressed more highly at the bud stage. (C) Gene Ontology biological-process enrichment analysis of genes expressed more highly in the combined 3-somite/6-somite population. For (B,C), bubble size indicates gene count, color represents *−* log_10_(*P* ), and the x-axis shows the enrichment factor.

## Supplementary Tables

**Table S1:**
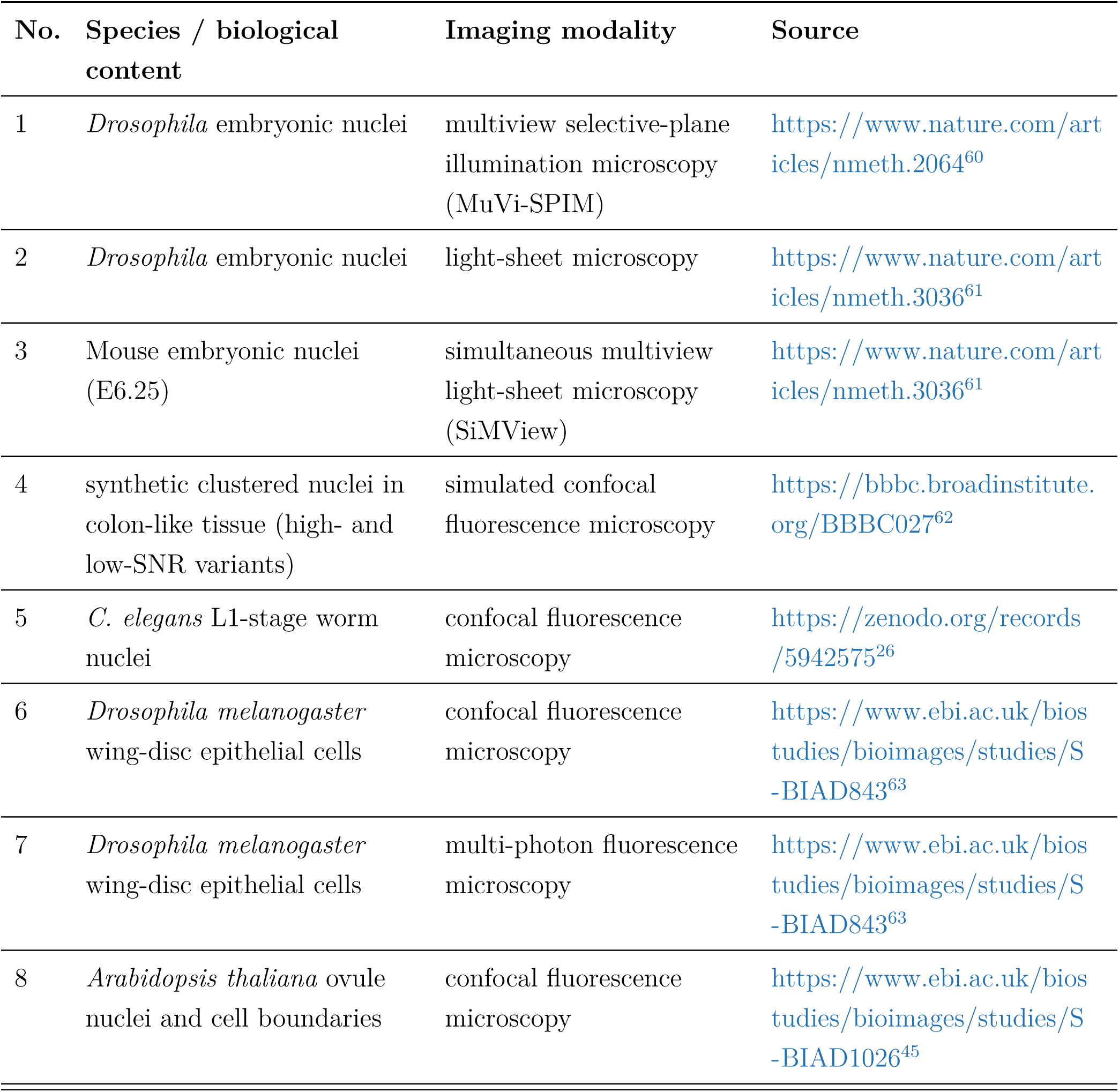
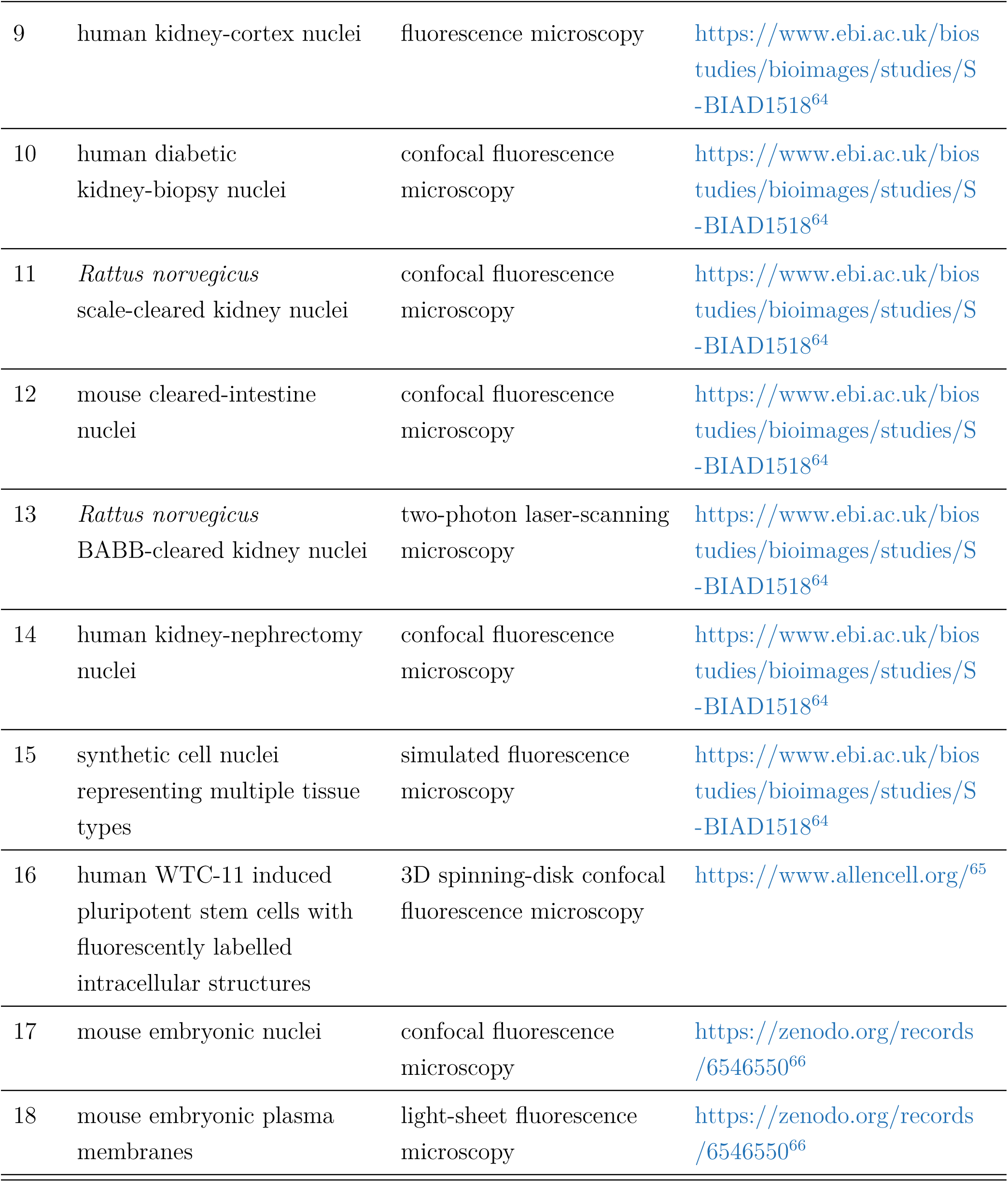
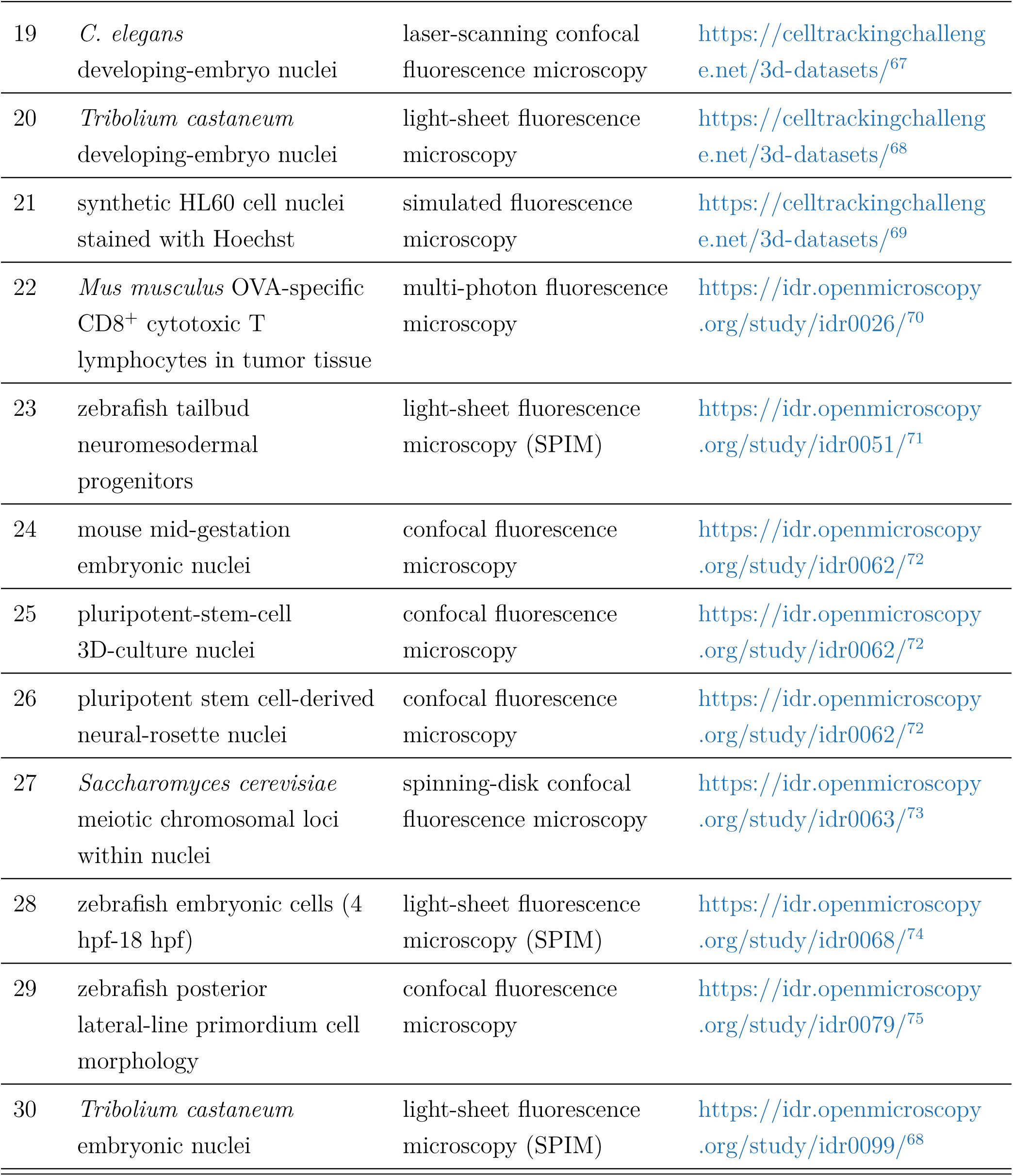
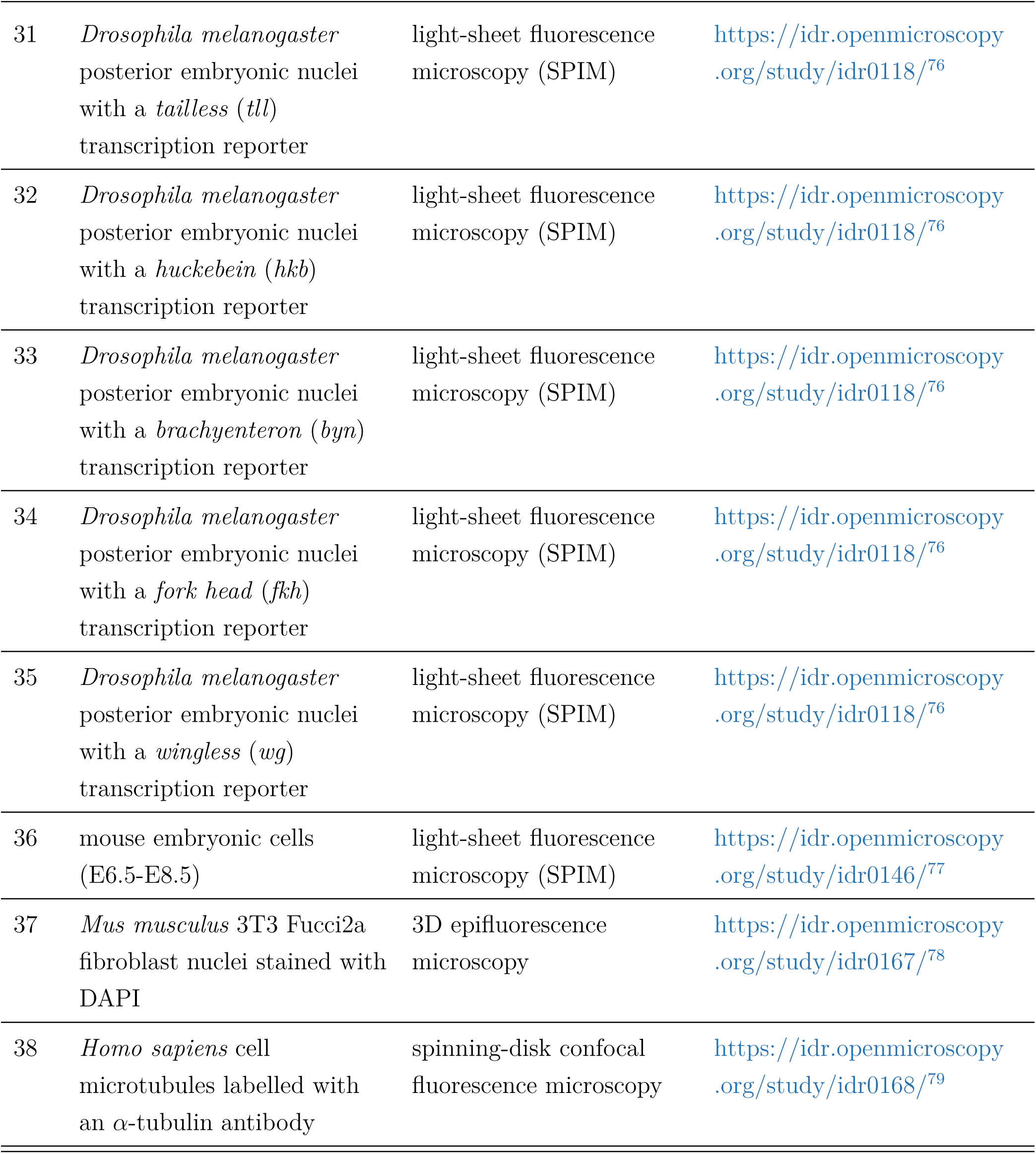
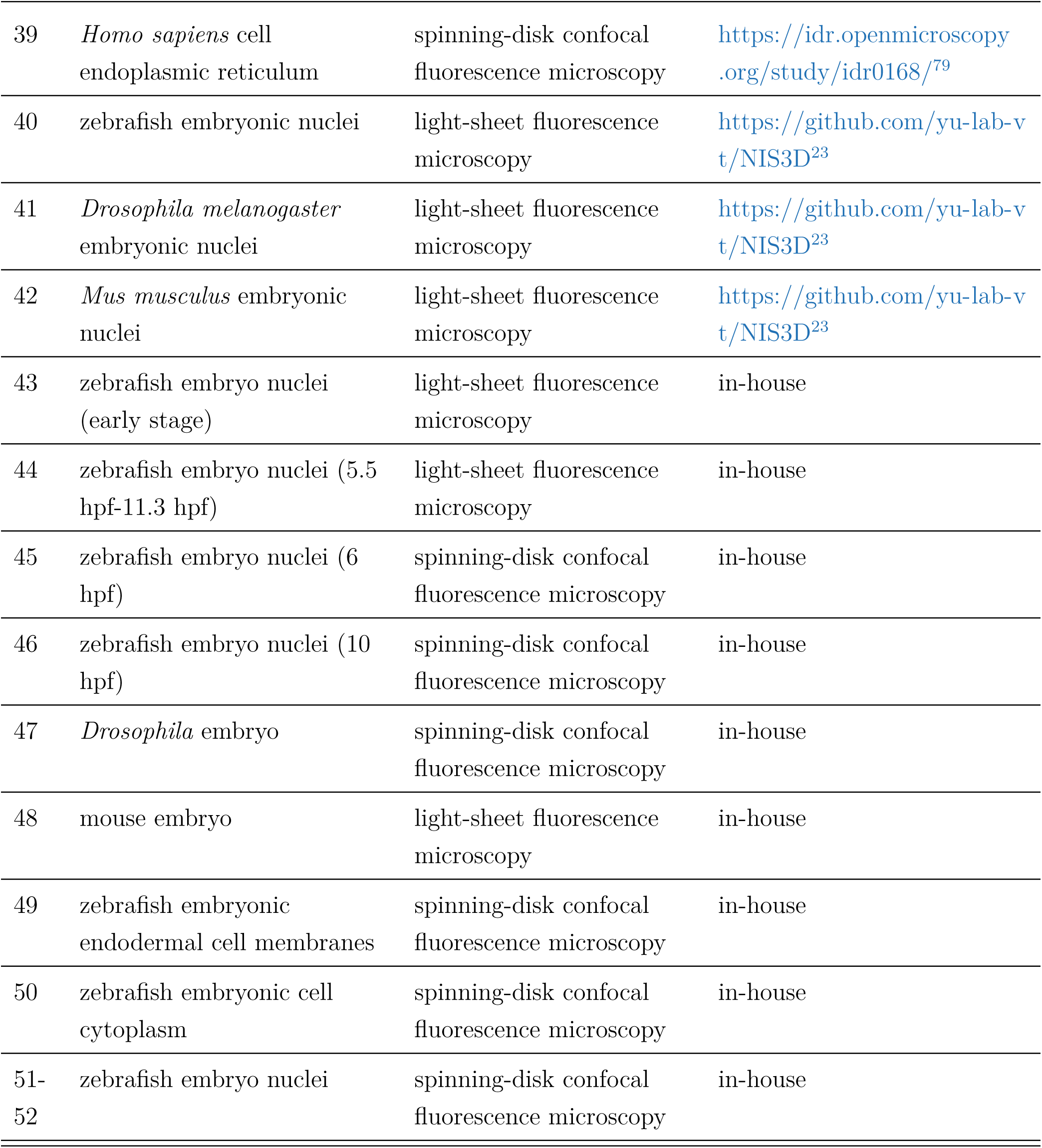
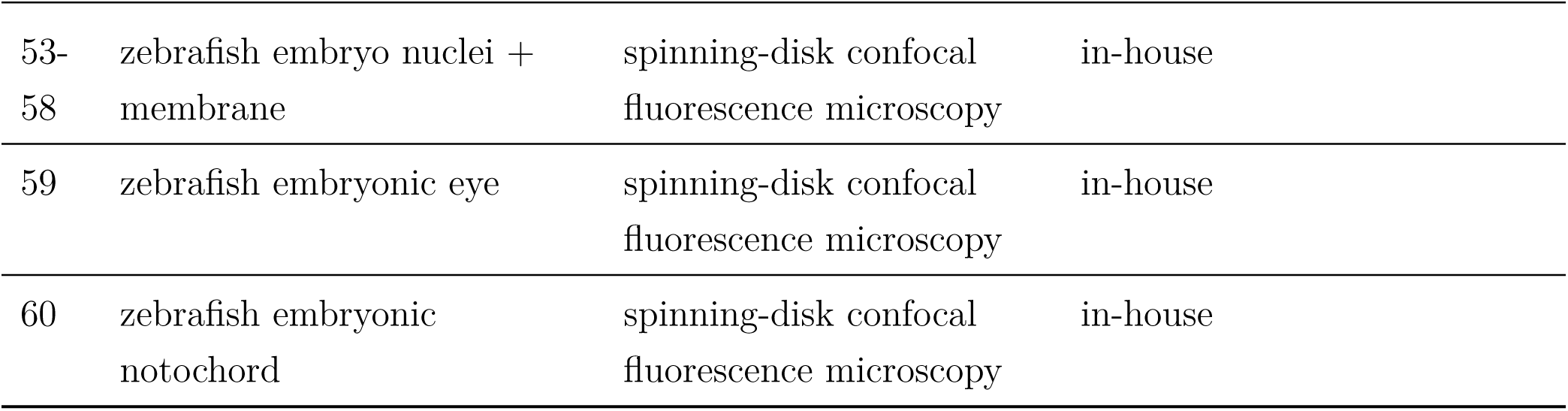
Summary of public and in-house 3D microscopy datasets used in this study. Different imaging modalities or content from the same dataset were listed separately in this table, resulting in 60 dataset–modality entries corresponding to the 47 datasets described in the main text.

| No. | Species / biological content | Imaging modality | Source |
| --- | --- | --- | --- |
| 1 | *Drosophila* embryonic nuclei | multiview selective-plane illumination microscopy (MuVi-SPIM) | https://www.nature.com/articles/nmeth.2064<sup>60</sup> |
| 2 | *Drosophila* embryonic nuclei | light-sheet microscopy | https://www.nature.com/articles/nmeth.3036<sup>61</sup> |
| 3 | Mouse embryonic nuclei (E6.25) | simultaneous multiview light-sheet microscopy (SiMView) | https://www.nature.com/articles/nmeth.3036<sup>61</sup> |
| 4 | synthetic clustered nuclei in colon-like tissue (high- and low-SNR variants) | simulated confocal fluorescence microscopy | https://bbbc.broadinstitute.org/BBBC027<sup>62</sup> |
| 5 | *C. elegans* L1-stage worm nuclei | confocal fluorescence microscopy | https://zenodo.org/records/5942575<sup>26</sup> |
| 6 | *Drosophila melanogaster* wing-disc epithelial cells | confocal fluorescence microscopy | https://www.ebi.ac.uk/biosudies/bioimages/studies/S-BIAD843<sup>63</sup> |
| 7 | *Drosophila melanogaster* wing-disc epithelial cells | multi-photon fluorescence microscopy | https://www.ebi.ac.uk/biosudies/bioimages/studies/S-BIAD843<sup>63</sup> |
| 8 | *Arabidopsis thaliana* ovule nuclei and cell boundaries | confocal fluorescence microscopy | https://www.ebi.ac.uk/biosudies/bioimages/studies/S-BIAD1026<sup>45</sup> |
| 9 | human kidney-cortex nuclei | fluorescence microscopy | https://www.ebi.ac.uk/biosudies/bioimages/studies/S-BIAD1518<sup>64</sup> |
| 10 | human diabetic kidney-biopsy nuclei | confocal fluorescence microscopy | https://www.ebi.ac.uk/biosudies/bioimages/studies/S-BIAD1518<sup>64</sup> |
| 11 | *Rattus norvegicus* scale-cleared kidney nuclei | confocal fluorescence microscopy | https://www.ebi.ac.uk/biosudies/bioimages/studies/S-BIAD1518<sup>64</sup> |
| 12 | mouse cleared-intestine nuclei | confocal fluorescence microscopy | https://www.ebi.ac.uk/biosudies/bioimages/studies/S-BIAD1518<sup>64</sup> |
| 13 | *Rattus norvegicus* BABB-cleared kidney nuclei | two-photon laser-scanning microscopy | https://www.ebi.ac.uk/biosudies/bioimages/studies/S-BIAD1518<sup>64</sup> |
| 14 | human kidney-nephrectomy nuclei | confocal fluorescence microscopy | https://www.ebi.ac.uk/biosudies/bioimages/studies/S-BIAD1518<sup>64</sup> |
| 15 | synthetic cell nuclei representing multiple tissue types | simulated fluorescence microscopy | https://www.ebi.ac.uk/biosudies/bioimages/studies/S-BIAD1518<sup>64</sup> |
| 16 | human WTC-11 induced pluripotent stem cells with fluorescently labelled intracellular structures | 3D spinning-disk confocal fluorescence microscopy | https://www.allencell.org/<sup>65</sup> |
| 17 | mouse embryonic nuclei | confocal fluorescence microscopy | https://zenodo.org/records/6546550<sup>66</sup> |
| 18 | mouse embryonic plasma membranes | light-sheet fluorescence microscopy | https://zenodo.org/records/6546550<sup>66</sup> |
| 19 | *C. elegans*<br>developing-embryo nuclei | laser-scanning confocal<br>fluorescence microscopy | https://celltrackingchallenge.net/3d-datasets/<sup>67</sup> |
| 20 | *Tribolium castaneum*<br>developing-embryo nuclei | light-sheet fluorescence<br>microscopy | https://celltrackingchallenge.net/3d-datasets/<sup>68</sup> |
| 21 | synthetic HL60 cell nuclei<br>stained with Hoechst | simulated fluorescence<br>microscopy | https://celltrackingchallenge.net/3d-datasets/<sup>69</sup> |
| 22 | *Mus musculus* OVA-specific<br>CD8<sup>+</sup> cytotoxic T<br>lymphocytes in tumor tissue | multi-photon fluorescence<br>microscopy | https://idr.openmicroscopy.org/study/idr0026/<sup>70</sup> |
| 23 | zebrafish tailbud<br>neuromesodermal<br>progenitors | light-sheet fluorescence<br>microscopy (SPIM) | https://idr.openmicroscopy.org/study/idr0051/<sup>71</sup> |
| 24 | mouse mid-gestation<br>embryonic nuclei | confocal fluorescence<br>microscopy | https://idr.openmicroscopy.org/study/idr0062/<sup>72</sup> |
| 25 | pluripotent-stem-cell<br>3D-culture nuclei | confocal fluorescence<br>microscopy | https://idr.openmicroscopy.org/study/idr0062/<sup>72</sup> |
| 26 | pluripotent stem cell-derived<br>neural-rosette nuclei | confocal fluorescence<br>microscopy | https://idr.openmicroscopy.org/study/idr0062/<sup>72</sup> |
| 27 | *Saccharomyces cerevisiae*<br>meiotic chromosomal loci<br>within nuclei | spinning-disk confocal<br>fluorescence microscopy | https://idr.openmicroscopy.org/study/idr0063/<sup>73</sup> |
| 28 | zebrafish embryonic cells (4<br>hpf-18 hpf) | light-sheet fluorescence<br>microscopy (SPIM) | https://idr.openmicroscopy.org/study/idr0068/<sup>74</sup> |
| 29 | zebrafish posterior<br>lateral-line primordium cell<br>morphology | confocal fluorescence<br>microscopy | https://idr.openmicroscopy.org/study/idr0079/<sup>75</sup> |
| 30 | *Tribolium castaneum*<br>embryonic nuclei | light-sheet fluorescence<br>microscopy (SPIM) | https://idr.openmicroscopy.org/study/idr0099/<sup>68</sup> |
| 31 | *Drosophila melanogaster* posterior embryonic nuclei with a *tailless* (*tll*) transcription reporter | light-sheet fluorescence microscopy (SPIM) | https://idr.openmicroscopy.org/study/idr0118/<sup>76</sup> |
| 32 | *Drosophila melanogaster* posterior embryonic nuclei with a *huckebein* (*hkb*) transcription reporter | light-sheet fluorescence microscopy (SPIM) | https://idr.openmicroscopy.org/study/idr0118/<sup>76</sup> |
| 33 | *Drosophila melanogaster* posterior embryonic nuclei with a *brachyenteron* (*byn*) transcription reporter | light-sheet fluorescence microscopy (SPIM) | https://idr.openmicroscopy.org/study/idr0118/<sup>76</sup> |
| 34 | *Drosophila melanogaster* posterior embryonic nuclei with a *fork head* (*fkh*) transcription reporter | light-sheet fluorescence microscopy (SPIM) | https://idr.openmicroscopy.org/study/idr0118/<sup>76</sup> |
| 35 | *Drosophila melanogaster* posterior embryonic nuclei with a *wingless* (*wg*) transcription reporter | light-sheet fluorescence microscopy (SPIM) | https://idr.openmicroscopy.org/study/idr0118/<sup>76</sup> |
| 36 | mouse embryonic cells (E6.5-E8.5) | light-sheet fluorescence microscopy (SPIM) | https://idr.openmicroscopy.org/study/idr0146/<sup>77</sup> |
| 37 | *Mus musculus* 3T3 Fucci2a fibroblast nuclei stained with DAPI | 3D epifluorescence microscopy | https://idr.openmicroscopy.org/study/idr0167/<sup>78</sup> |
| 38 | *Homo sapiens* cell microtubules labelled with an α-tubulin antibody | spinning-disk confocal fluorescence microscopy | https://idr.openmicroscopy.org/study/idr0168/<sup>79</sup> |
| 39 | *Homo sapiens* cell endoplasmic reticulum | spinning-disk confocal fluorescence microscopy | https://idr.openmicroscopy.org/study/idr0168/<sup>79</sup> |
| 40 | zebrafish embryonic nuclei | light-sheet fluorescence microscopy | https://github.com/yu-lab-vt/NIS3D<sup>23</sup> |
| 41 | *Drosophila melanogaster* embryonic nuclei | light-sheet fluorescence microscopy | https://github.com/yu-lab-vt/NIS3D<sup>23</sup> |
| 42 | *Mus musculus* embryonic nuclei | light-sheet fluorescence microscopy | https://github.com/yu-lab-vt/NIS3D<sup>23</sup> |
| 43 | zebrafish embryo nuclei (early stage) | light-sheet fluorescence microscopy | in-house |
| 44 | zebrafish embryo nuclei (5.5 hpf-11.3 hpf) | light-sheet fluorescence microscopy | in-house |
| 45 | zebrafish embryo nuclei (6 hpf) | spinning-disk confocal fluorescence microscopy | in-house |
| 46 | zebrafish embryo nuclei (10 hpf) | spinning-disk confocal fluorescence microscopy | in-house |
| 47 | *Drosophila* embryo | spinning-disk confocal fluorescence microscopy | in-house |
| 48 | mouse embryo | light-sheet fluorescence microscopy | in-house |
| 49 | zebrafish embryonic endodermal cell membranes | spinning-disk confocal fluorescence microscopy | in-house |
| 50 | zebrafish embryonic cell cytoplasm | spinning-disk confocal fluorescence microscopy | in-house |
| 51-52 | zebrafish embryo nuclei | spinning-disk confocal fluorescence microscopy | in-house |
| 53-58 | zebrafish embryo nuclei + membrane | spinning-disk confocal fluorescence microscopy | in-house |
| 59 | zebrafish embryonic eye | spinning-disk confocal fluorescence microscopy | in-house |
| 60 | zebrafish embryonic notochord | spinning-disk confocal fluorescence microscopy | in-house |

**Table S2:** Summary of gold and silver annotations in public datasets.

| No. | Species / content | Annotation type | Annotation source |
| --- | --- | --- | --- |
| 6,7 | *Drosophila* wing disc | Gold | Public |
| 8 | Arabidopsis ovule nuclei and cell boundaries | Gold | Public |
| 17 | mouse embryonic nuclei | Gold | Public |
| 40-42 | 3D nuclei | Gold | Public |
| 1 | *Drosophila* embryo | Silver | PrinCut-Auto |
| 2 | *Drosophila* embryo | Silver | PrinCut-Auto |
| 4 | synthetic colon tissue | Silver | Public |
| 9-14 | human/rat/mouse nuclei | Silver | Public |
| 15 | synthetic cell nuclei | Silver | Public |
| 18 | mouse embryo membrane | Silver | Public |
| 20 | *Tribolium* embryo | Silver | PrinCut-Auto |
| 21 | simulated HL60 nuclei | Silver | Public |
| 29 | zebrafish lateral line | Silver | Public |
| 31-35 | *Drosophila* endoderm | Silver | Public |

**Table S3:** Summary of FOCUS-3D gold annotations.

| No. | Species / content | Number of images | Number of cells | Data source |
| --- | --- | --- | --- | --- |
| 1 | *Drosophila* embryo | 5 | 3,713 | Public |
| 2 | *Drosophila* embryo | 9 | 74,630 | Public |
| 3 | mouse embryo | 5 | 45,531 | Public |
| 16 | human WTC-11 iPS cells | 5 | 219 | Public |
| 19 | *C. elegans* | 10 | 1,404 | Public |
| 43 | zebrafish embryo nuclei | 10 | 76,114 | In-house |
| 44 | zebrafish embryo nuclei | 10 | 181,771 | In-house |
| 53 | zebrafish embryo nuclei | 4 | 6,931 | In-house |
| 59,60 | zebrafish embryo nuclei | 5 | 17,738 | In-house |
| 53 | zebrafish embryo membrane | 6 | 3,552 | In-house |
| 49 | zebrafish embryo membrane | 3 | 2,233 | In-house |
| 50 | zebrafish embryo membrane<br>+ cytoplasm | 10 | 382 | In-house |

## References

[1] Katie McDole, et al. “In toto imaging and reconstruction of post-implantation mouse development at the single-cell level”. In: Cell 175.3 (2018), pp. 859–876.

[2] Hans Clevers. “Modeling development and disease with organoids”. In: Cell 165.7 (2016), pp. 1586–1597.

[3] Erick Moen, et al. “Deep learning for cellular image analysis”. In: Nature Methods 16.12 (2019), pp. 1233–1246.

[4] Erik Meijering. “Cell segmentation: 50 years down the road [life sciences]”. In: IEEE Signal Processing Magazine 29.5 (2012), pp. 140–145.

[5] Johannes Stegmaier, et al. “Real-time three-dimensional cell segmentation in large-scale microscopy data of developing embryos”. In: Developmental Cell 36.2 (2016), pp. 225–240.

[6] Noah F Greenwald, et al. “Whole-cell segmentation of tissue images with human-level performance using large-scale data annotation and deep learning”. In: Nature Biotechnology 40.4 (2022), pp. 555–565.

[7] Vladimír Ulman, et al. “An objective comparison of cell-tracking algorithms”. In: Nature Methods 14.12 (2017), pp. 1141–1152.

[8] Mengfan Wang, et al. “High-Fidelity Long-term Whole-embryo Lineage and Fate Reconstruction by Iterative Tracking with Error Correction”. In: bioRxiv (2026).

[9] Özgün Çiçek et al. “3D U-Net: learning dense volumetric segmentation from sparse annotation”. In: International conference on medical image computing and computer-assisted intervention. Springer. 2016, pp. 424–432.

[10] Ashish Vaswani, et al. “Attention is all you need”. In: Advances in Neural Information Processing Systems 30 (2017).

[11] Alexey Dosovitskiy, et al. “An image is worth 16×16 words: Transformers for image recognition at scale”. In: arXiv preprint arXiv:2010.11929 (2020).

[12] Adrian Wolny, et al. “Accurate and versatile 3D segmentation of plant tissues at cellular resolution”. In: eLife 9 (2020), e57613.

[13] Carsen Stringer, et al. “Cellpose: a generalist algorithm for cellular segmentation”. In: Nature Methods 18.1 (2021), pp. 100–106.

[14] Kevin J Cutler, et al. “Omnipose: a high-precision morphology-independent solution for bacterial cell segmentation”. In: Nature Methods 19.11 (2022), pp. 1438–1448.

[15] Chentao Wen, et al. “Seg2Link: an efficient and versatile solution for semi-automatic cell segmentation in 3D image stacks”. In: Scientific Reports 13.1 (2023), p. 7109.

[16] Markus Marks, et al. “CellSAM: a foundation model for cell segmentation”. In: Nature Methods 22 (2025), pp. 2585–2593.

[17] Haoran Chen, et al. “3DCellComposer - A versatile pipeline utilizing 2D cell segmentation methods for 3D cell segmentation”. In: Methods (2025).

[18] Danny Salem, et al. “Yeastnet: Deep-learning-enabled accurate segmentation of budding yeast cells in bright-field microscopy”. In: Applied Sciences 11.6 (2021), p. 2692.

[19] Juan C Caicedo, et al. “Nucleus segmentation across imaging experiments: the 2018 Data Science Bowl”. In: Nature Methods 16.12 (2019), pp. 1247–1253.

[20] Fabian Hörst, et al. “CellViT: Vision transformers for precise cell segmentation and classification”. In: Medical Image Analysis 94 (2024), p. 103143.

[21] Andong Wang, et al. “A novel deep learning-based 3D cell segmentation framework for future image-based disease detection”. In: Scientific Reports 12.1 (2022), p. 342.

[22] Liming Wu, et al. “NISNET3D: Three-dimensional nuclear synthesis and instance segmentation for fluorescence microscopy images”. In: Scientific Reports 13.1 (2023), p. 9533.

[23] Wei Zheng, et al. “NIS3D: a completely annotated benchmark for dense 3D nuclei image segmentation”. In: Advances in Neural Information Processing Systems 36 (2023), pp. 4741– 4752.

[24] Cyril Achard, et al. “CellSeg3D, Self-supervised 3D cell segmentation for fluorescence microscopy”. In: eLife 13 (2025), RP99848.

[25] Dennis Eschweiler, Richard S Smith, and Johannes Stegmaier. “Robust 3D cell segmentation: extending the view of cellpose”. In: 2022 IEEE International Conference on Image Processing (ICIP). IEEE. 2022, pp. 191–195.

[26] Martin Weigert et al. “Star-convex polyhedra for 3D object detection and segmentation in microscopy”. In: Proceedings of the IEEE/CVF winter conference on applications of computer vision. 2020, pp. 3666–3673.

[27] Felix Y Zhou, et al. “Universal consensus 3D segmentation of cells from 2D segmented stacks”. In: Nature Methods 22 (2025), pp. 2386–2399.

[28] Anwai Archit, et al. “Segment anything for microscopy”. In: Nature Methods 22.3 (2025), pp. 579–591.

[29] Bowen Cheng et al. “Masked-attention mask transformer for universal image segmentation”. In: Proceedings of the IEEE/CVF conference on computer vision and pattern recognition. 2022, pp. 1290–1299.

[30] Feng Li et al. “Mask DINO: Towards a unified transformer-based framework for object detection and segmentation”. In: Proceedings of the IEEE/CVF conference on computer vision and pattern recognition. 2023, pp. 3041–3050.

[31] Marius Pachitariu and Carsen Stringer. “Cellpose 2.0: how to train your own model”. In: Nature Methods 19.12 (2022), pp. 1634–1641.

[32] Stuart Berg, et al. “Ilastik: interactive machine learning for (bio) image analysis”. In: Nature Methods 16.12 (2019), pp. 1226–1232.

[33] Toby GR Andrews, et al. “Single-cell morphometrics reveals ancestral principles of notochord development”. In: Development 148.16 (2021), dev199430.

[34] Yinan Wan, et al. “Whole-embryo spatial transcriptomics at subcellular resolution from gastrulation to organogenesis”. In: Science 391.6790 (2026), eadt3439.

[35] Jeffrey A Farrell, et al. “Single-cell reconstruction of developmental trajectories during zebrafish embryogenesis”. In: Science 360.6392 (2018), eaar3131.

[36] Wei Zheng et al. “PrinCut-Auto: An Unsupervised 3D Cell Detection Tool for Embryonic Data”. In: 2024 IEEE International Symposium on Biomedical Imaging (ISBI). IEEE. 2024, pp. 1–5.

[37] Kaiming He et al. “Masked autoencoders are scalable vision learners”. In: Proceedings of the IEEE/CVF conference on computer vision and pattern recognition. 2022, pp. 16000–16009.

[38] Jacob Devlin et al. “BERT: Pre-training of Deep Bidirectional Transformers for Language Understanding”. In: Proceedings of the 2019 Conference of the North American Chapter of the Association for Computational Linguistics: Human Language Technologies. 2019, pp. 4171–4186.

[39] Tom B. Brown, et al. “Language Models are Few-Shot Learners”. In: Advances in Neural Information Processing Systems 33 (2020), pp. 1877–1901.

[40] Hugo Touvron, et al. “Llama 2: Open Foundation and Fine-Tuned Chat Models”. In: arXiv preprint arXiv:2307.09288 (2023).

[41] Zhe Chen, et al. “Vision transformer adapter for dense predictions”. In: arXiv preprint arXiv:2205.08534 (2022).

[42] Hao Zhang, et al. “DINO: DETR with improved denoising anchor boxes for end-to-end object detection”. In: arXiv preprint arXiv:2203.03605 (2022).

[43] Yingming Wang et al. “Anchor DETR: Query design for transformer-based detector”. In: Proceedings of the AAAI conference on artificial intelligence. Vol. 36. 2022, pp. 2567–2575.

[44] Marius Pachitariu, Michael Rariden, and Carsen Stringer. “Cellpose-SAM: superhuman generalization for cellular segmentation”. In: bioRxiv (2025). doi: 10.1101/2025.04.28.651001.

[45] Athul Vijayan, et al. “A deep learning-based toolkit for 3D nuclei segmentation and quantitative analysis in cellular and tissue context”. In: Development 151.14 (2024), dev202800.

[46] napari contributors. napari: a multi-dimensional image viewer for Python. 2019. doi: 10.5281/zenodo.3555620. url: https://doi.org/10.5281/zenodo.3555620.

[47] Masazumi Tada and Carl-Philipp Heisenberg. “Convergent extension: using collective cell migration and cell intercalation to shape embryos”. In: Development 139.21 (2012), pp. 3897– 3904.

[48] Jordão Bragantini, et al. “Ultrack: pushing the limits of cell tracking across biological scales”. In: Nature Methods 22.11 (2025), pp. 2423–2436.

[49] Nathalia S. Glickman et al. “Shaping the zebrafish notochord”. In: Development 130.5 (2003), pp. 873–887. doi: 10.1242/dev.00314.

[50] Fang Lin et al. “Essential roles of G*α*12/13 signaling in distinct cell behaviors driving zebrafish convergence and extension gastrulation movements”. In: The Journal of Cell Biology 169.5 (2005), pp. 777–787.

[51] Margot L. K. Williams et al. “Gon4l regulates notochord boundary formation and cell polarity underlying axis extension by repressing adhesion genes”. In: Nature Communications 9.1 (2018), p. 1319. doi: 10.1038/s41467-018-03715-w.

[52] Merlin Lange, et al. “A multimodal zebrafish developmental atlas reveals the state-transition dynamics of late-vertebrate pluripotent axial progenitors”. In: Cell 187.23 (2024), pp. 6742– 6759.

[53] Denise Serra, et al. “Self-organization and symmetry breaking in intestinal organoid development”. In: Nature 569.7754 (2019), pp. 66–72.

[54] Zhisong He, et al. “Lineage recording in human cerebral organoids”. In: Nature Methods 19.1 (2022), pp. 90–99.

[55] Çağrı Çevrim, et al. “Long-term live imaging, cell identification and cell tracking in regenerating crustacean legs”. In: eLife 14 (2025), RP107534.

[56] Yiqun Wang, et al. “Gene module reconstruction identifies cellular differentiation processes and the regulatory logic of specialized secretion in zebrafish”. In: Developmental Cell 60.4 (2025), pp. 581–598.

[57] Ricardo Londono, et al. “Tissue repair and epimorphic regeneration: an overview”. In: Current Pathobiology Reports 6.1 (2018), pp. 61–69.

[58] Sungmin Baek, et al. “Single-cell transcriptome analysis reveals three sequential phases of gene expression during zebrafish sensory hair cell regeneration”. In: Developmental Cell 57.6 (2022), pp. 799–819.

[59] Valentina Cigliola, Clayton J Becker, and Kenneth D Poss. “Building bridges, not walls: spinal cord regeneration in zebrafish”. In: Disease Models & Mechanisms 13.5 (2020), p. dmm044131.

## Supplementary References

[60] Uros Krzic, et al. “Multiview light-sheet microscope for rapid in toto imaging”. In: Nature Methods 9.7 (2012), pp. 730–733.

[61] Fernando Amat, et al. “Fast, accurate reconstruction of cell lineages from large-scale fluorescence microscopy data”. In: Nature Methods 11.9 (2014), pp. 951–958.

[62] David Svoboda, Ondøej Homola, and Stanislav Stejskal. “Generation of 3D digital phantoms of colon tissue”. In: International Conference Image Analysis and Recognition. Springer. 2011, pp. 31–39.

[63] Giulia Paci, et al. “Single cell resolution 3D imaging and segmentation within intact live tissues”. In: npj Imaging 3.1 (2025), p. 40.

[64] Alain Chen, et al. “3D ground truth annotations of nuclei in 3D microscopy volumes”. In: bioRxiv (2022).

[65] Matheus P Viana, et al. “Integrated intracellular organization and its variations in human iPS cells”. In: Nature 613.7943 (2023), pp. 345–354.

[66] Vladyslav Bondarenko, et al. “Embryo-uterine interaction coordinates mouse embryogenesis during implantation”. In: The EMBO Journal 42.17 (2023), e113280.

[67] John Isaac Murray, et al. “Automated analysis of embryonic gene expression with cellular resolution in C. elegans”. In: Nature Methods 5.8 (2008), pp. 703–709.

[68] Akanksha Jain, et al. “Regionalized tissue fluidization is required for epithelial gap closure during insect gastrulation”. In: Nature Communications 11.1 (2020), p. 5604.

[69] David Svoboda and Vladimir Ulman. “MitoGen: a framework for generating 3D synthetic time-lapse sequences of cell populations in fluorescence microscopy”. In: IEEE Transactions on Medical Imaging 36.1 (2016), pp. 310–321.

[70] Bettina Weigelin, et al. “Focusing and sustaining the antitumor CTL effector killer response by agonist anti-CD137 mAb”. In: Proceedings of the National Academy of Sciences 112.24 (2015), pp. 7551–7556.

[71] Andrea Attardi, et al. “Neuromesodermal progenitors are a conserved source of spinal cord with divergent growth dynamics”. In: Development 145.21 (2018), dev166728.

[72] Guillaume Blin, et al. “Nessys: a new set of tools for the automated detection of nuclei within intact tissues and dense 3D cultures”. In: PLoS Biology 17.8 (2019), e3000388.

[73] Trent AC Newman, et al. “Diffusion and distal linkages govern interchromosomal dynamics during meiotic prophase”. In: Proceedings of the National Academy of Sciences 119.12 (2022), e2115883119.

[74] Gopi Shah, et al. “Multi-scale imaging and analysis identify pan-embryo cell dynamics of germlayer formation in zebrafish”. In: Nature Communications 10.1 (2019), p. 5753.

[75] Jonas Hartmann, et al. “An image-based data-driven analysis of cellular architecture in a developing tissue”. In: eLife 9 (2020), e55913.

[76] Shannon E Keenan, et al. “Dynamics of Drosophila endoderm specification”. In: Proceedings of the National Academy of Sciences 119.15 (2022), e2112892119.

[77] Martin H Dominguez, et al. “Graded mesoderm assembly governs cell fate and morphogenesis of the early mammalian heart”. In: Cell 186.3 (2023), pp. 479–496.

[78] Gang Li, et al. “Predicting cell cycle stage from 3D single-cell nuclear-stained images”. In: Life Science Alliance 8.6 (2025).

[79] Xinyi Zhang, et al. “Prediction of protein subcellular localization in single cells”. In: Nature Methods 22.6 (2025), pp. 1265–1275.

